# DensityPath: a level-set algorithm to visualize and reconstruct cell developmental trajectories for large-scale single-cell RNAseq data

**DOI:** 10.1101/276311

**Authors:** Ziwei Chen, Shaokun An, Xiangqi Bai, Fuzhou Gong, Liang Ma, Lin Wan

## Abstract

Cell fates are determined by transition-states which occur during complex biological processes such as proliferation and differentiation. The advance in single-cell RNA sequencing (scRNAseq) provides the snapshots of single cell transcriptomes, thus offering an essential opportunity to study such complex biological processes. Here, we introduce a novel algorithm, DensityPath, which visualizes and reconstructs the underlying cell developmental trajectories for large-scale scRNAseq data. DensityPath has three merits. Firstly, by adopting the nonlinear dimension reduction algorithm elastic embedding, DensityPath reveals the intrinsic structures of the data. Secondly, by applying the powerful level set clustering method, DensityPath extracts the separate high density clusters of representative cell states (RCSs) from the single cell multimodal density landscape of gene expression space, enabling it to handle the heterogeneous scRNAseq data elegantly and accurately. Thirdly, DensityPath constructs cell state-transition path by finding the geodesic minimum spanning tree of the RCSs on the surface of the density landscape, making it more computationally efficient and accurate for large-scale dataset. The cell state-transition path constructed by DensityPath has the physical interpretation as the minimum-transition-energy (least-cost) path. We demonstrate that DensityPath is capable of identifying complex cell development trajectories with bifurcating and trifurcating branches on the human preimplantation embryos. We demonstrate that DensityPath is robust and has high accuracy of pseudotime calculation and branch assignment on the real scRNAseq as well as simulated datasets.

## Introduction

Cell fates are determined by transition-states which occur during complex biological processes such as proliferation and differentiation. The understanding of the mechanism of cell fate decisions is one of the most fundamental questions in biology [1]. The recent advent of massively parallel single-cell RNA sequencing (scRNAseq) provides the snapshots of single cell transcriptomes, thus offering an essential opportunity to unveil the molecular mechanism of cell fate decisions [2, 3]. The single cells assayed are undergoing asynchronous processes, while temporal and spatial information of each cell is generally unavailable. Thus, the cellular profiles by scRNAseq are heterogeneous populations of cells at diverse states of cell proliferation and differentiation, making it a computational challenge to infer the progression of cell fate decisions based on scRNAseq data. Here we present a novel algorithm, DensityPath, which accurately and efficiently visualizes and reconstructs the underlying cell developmental trajectories for large-scale scRNAseq data. DensityPath, which is based on the powerful mathematical tool of level set clustering [4, 5, 6], handles the heterogeneity of scRNAseq data elegantly and accurately, addressing a critical problem of the study of epigenetic landscapes.

Intensive efforts have been devoted to computationally reconstructing pseudo trajectories of cellular development from single cell data (see [7, 8] for recent reviews). Monocle, which relied on building the minimum spanning tree (MST) on the cells, pioneered the study of reconstruction of complex pseudo trajectory for scRNAseq data [2]. However, Monocle is computationally intractable for large-scale data. Wanderlust, another pioneer study, attempted to construct a linear trajectory based on the *k*-nearest neighbor (*k*-NN) graph for the single cell mass cytometry data [9]. To further reveal the hierarchical structure of cell lineage, Wishbone extended the Wanderlust algorithm to construct developmental bifurcating trajectory with two branches [10]. Diffusion pseudotime (DPT) developed a diffusion distance to calculate the diffusion pseudotime to reconstruct the branched trajectory [11]. Although it is possible to identify the lineage bifurcating event(s) for large-scale single cell data, Wishbone and DPT only construct simple branched trajectories, restricting to two branches or requiring the number of branches as *a prior*.

In order to reconstruct complex cell developmental trajectories at affordable computational cost, methods based on the embedding curve/graph techniques have been developed [12, 13, 14, 15]. These methods fitted/mapped the *d*-dimensional single cell data points onto one-dimensional curve or graph, and then ordered the cells along the curve/graph to approximate cell developmental trajectory. SCUBA [12] fitted the single cell data points with the principal curve, a smooth one-dimensional curve that passes through the “middle” data points [16, 17]. But due to the necessity of being infinitely differentiable, principal curve cannot handle self-intersecting data structure (e.g. the branched trajectory) [18]. Thus, SCUBA had to require additional temporal information and conduct the *K*-means algorithm on the temporal windows to extract cellular lineage relationships. TSCAN [13] also used a cluster-based approach to group cells into clusters by the hierarchical clustering algorithm, construct a MST by connecting cluster centers, and then project each cell onto the tree to obtain the pseudotime of cells. ScTDA [14] applied the topological data analysis (TDA) tool called Mapper [19] to the single cell data, intending to unveil the intrinsic topological structures of cell developmental processes. The Reeb graph constructed by scTDA includes not only tree-like structures, but also loops or holes. However, these TDA methods may be sensitive to noises of data [18]. Monocle2 [15], a descendant of Monocle, adapted a recently developed principal graph method, reversed graph embedding (RGE) [18], to find the complex trajectories of single cell data. Principal graph handles the self-intersecting data structure by a collection of piecewise smooth curves where these curves can intersect each other [18]. To implement it, RGE utilized *K*-means algorithm to obtain *K* centroids of clusters, and then found the spanning tree between the centroids to construct the principal graph. RGE outperformed Mapper in constructing the progression path of breast cancer data [18].

Single cell gene expression profiles by scRNAseq are of high-dimensionality. Therefore, in most algorithms for constructing cell developmental trajectories, the very first and indeed a key step is to embed the single cell expression data onto a lower-dimensional space. The dimension-reduction and visualization methods such as principal component analysis (PCA), diffusion map, and t-distributed Stochastic Neighborhood Embedding (tSNE) are commonly used for scRNAseq (see [7, 8] for recent reviews). However, as pointed out by Moon et al [20] these methods have limitations to recover the complex structures of the scRNAseq data: PCA is a linear method which cannot handle the nonlinear structures; diffusion map can learn nonlinear transition paths but tends to place different branches into different diffusion dimensions, while tSNE can reveal and emphasize the cluster structures in data but tends to shatter trajectories and cannot preserve the global structures. Recently, a newly proposed extension of tSNE, elastic embedding (EE) [21], which exhibits the potentiality of visualizing the intrinsic latent structures of the scRNAseq data with both global and local manifold structures preserved, while improves the robustness and computational efficiency.

In this study, we propose a new algorithm, DensityPath, to reconstruct complex cell developmental trajectories for large-scale single cell data with an alternative strategy to find the embedding trajectory. Instead of fitting data to the principal graph, DensityPath first estimates the single cell density landscape on the embedded space, which conducted by EE, of gene expression profile.

Then, by level set method [4], DensityPath analyzes the heterogeneous multimodal behavior of the density landscape to identify the representative cell states (RCSs) and extract high density clusters. The level set method is a mathematical tool for the numerical analysis of surfaces and shapes, which has successful applications in unsupervised clustering, image processing, computational fluid dynamics, etc. [4, 6, 22, 23]. The intuition behind it is to reduce overall data complexity by breaking complex dataset into a series of separate high-density clusters, each of which is then regarded as a representative class of data [24].

Finally, DensityPath constructs the cell state-transition path by calculating the shortest path (geodesic), based on RCSs, of cell states on the surface of single cell density landscape and finding their MST. In order to connect these RCSs, we find that existing metrics on the level set clusters mainly consider the “height” at which two points or two clusters merge on the density function [22], which is insufficient to measure the transition distance between the points or clusters on the density function. Therefore, in DensityPath, we propose a metric of differential geometry for the RCSs. In detail, instead of using the Euclidean distance or density tree metric, DensityPath calculates the geodesic distances (shortest path distance) of the single cell points on the surface of single cell density landscape. By regarding the peak points of the RCSs identified as the landmarks, DensityPath reconstructs the cell state-transition path by finding the MST of the peak points based on their calculated geodesic distances. It is worth noting that the energy function can be defined as the negative logarithm of the density [25, 26]. It is therefore, the cell state-transition path based on the geodesic distance constructed by DensityPath exhibit physical interpretation as the shortest path, which is the minimum-transition-energy (least-cost) path, on the cell state energy landscape.

An overview of the procedures applied by DensityPath is given in Figure 1. We will apply DensityPath to real scRNAseq data of the mouse bone marrow cells [27], mouse hematopoietic stem and progenitor cells (HSPCs) bifurcating to myeloid and erythroid precursors [10], human preimplantation embryos [28], as well as two simulated datasets from [20, 29]. The performance in construction of cell developmental trajectory as well as the accuracy of pseudotime calculation and branch assignment of DensityPath were compared to those of existing methods.

**Figure 1:**
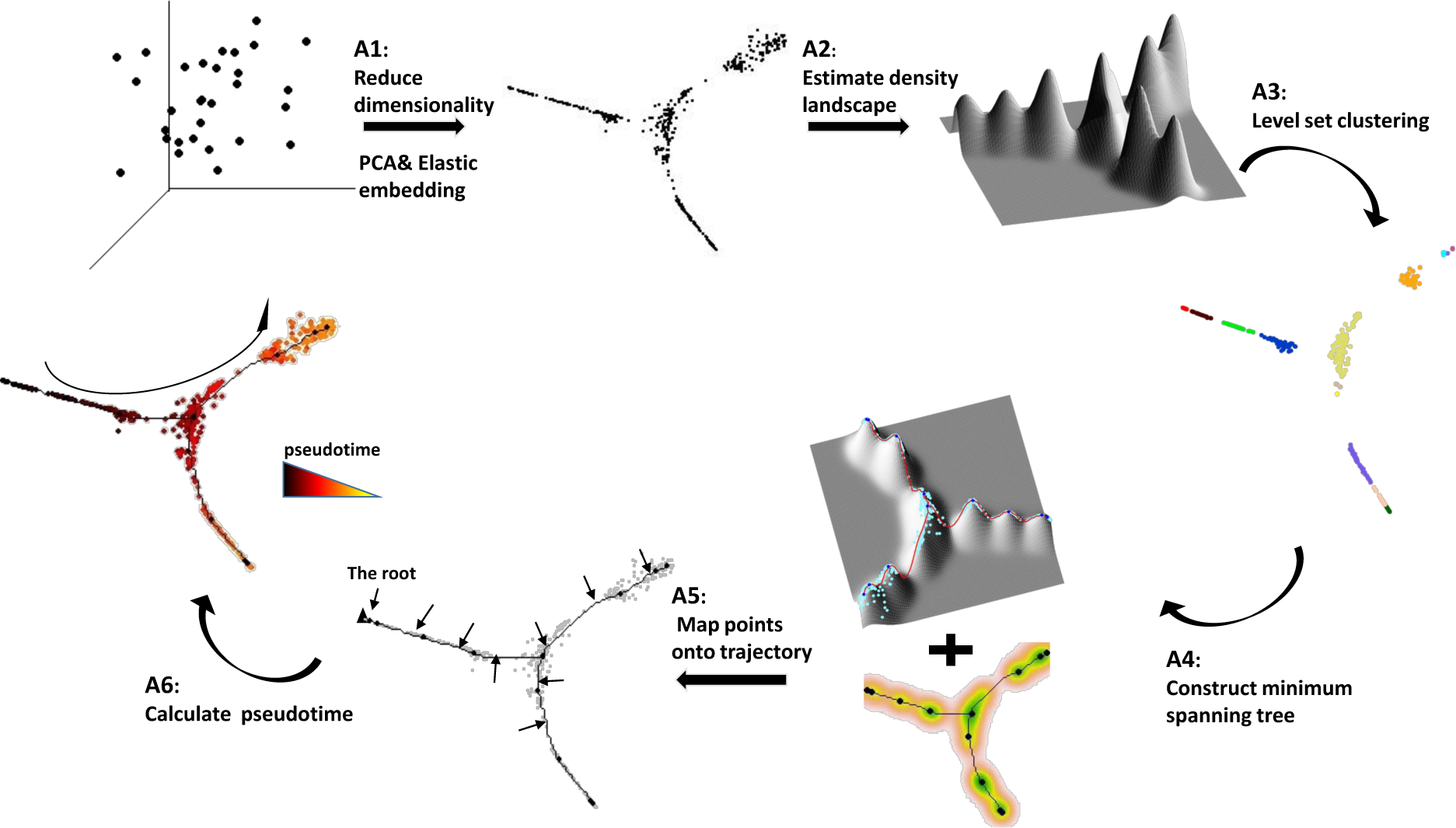
Overview of DensityPath. DensityPath reconstructs the trajectory of a differentiation progression and calculates the pseudotime of each cell along the path robustly and efficiently. Each cell is represented as a point in high-dimensional space, where each dimension corresponds to the expression level of a gene. Data are projected onto a 2-dimensional space by dimension reduction technique PCA and EE, and a density landscape is constructed through KDE method. Meanwhile, level set clustering method is applied to extract RCSs automatically and a minimum spanning tree is constructed on the surface of density landscape, connecting the density peaks of each RCSs. To each cell, DensityPath finds its nearest point on the trajectory, which has the smallest geodesic on the surface of the landscape. If a start cell (the root) is given, the pseudotime of each cell can be calculated.

## Methods

### Nonlinear dimension reduction algorithm elastic embedding

For a collection of data points *y*_1_, *…, y*_*n*_ ∈ ℝ^*d*^ in high dimensionality, elastic embedding (EE) algorithm [21] learns the latent *d*-dimensional coordinates *X*_*d×n*_ = (*x*_1_, *…, x*_*n*_) by optimizing the following objective function

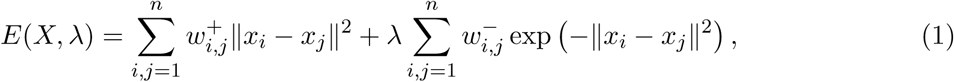

where *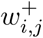* is the attractive weight (e.g. Gaussian affinity) between the data points, 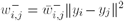 is the repulsive weight, and *λ ≥* 0 is the regularization parameter. By penalizing placing far apart latent points from similar data points and placing close together latent points from dissimilar data points, EE learns both the latent coordinates and the affinities between data points [21]. Compared with other techniques, EE preserves global and local manifold structure [6]. EE is efficient to large scale datasets, up to millions of points, by using spectral methods to increase computational speed [30]. Since the sample sizes of the datasets analyzed in this study are all *<* 5000, we adopted the EE algorithm of [31] to compute exact gradients, which is very efficient for data with sample size up to thousands.

### Level set clustering

We give a conceptual introduction to level sets and a level set clustering method called density tree. More details can be found in recent papers [6, 22].

Suppose we have a collection of samples *x*_1_, *…, x*_*n*_ *∈ℝ*^*d*^ that are independently drawn from an unknown probability distribution *F* with probability density function *f*. For any *t ≥* 0, the *t*-upper level set is defined as

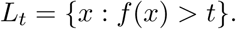

The connected components of *L*_*t*_, denoted as *𝓒*_*t*_, are called the density clusters at level *t*. Figure S1(a) illustrates that for a given 2-d density function, as *t* increasing, the size of *L*_*t*_ will decrease; after *t* going up across the valley of the adjacent peaks, a connected component of *L*_*t*_ will be broken into two connected components; after *t* going up across a peak, one connected component will vanish; after *t* going up across the highest peak of the density, *L*_*t*_ will be empty. By varying the threshold *t*, level set method thus analyzes the heterogeneous multimodal density landscape by examining how density cluster *𝓒*_*t*_ behaves on the density function.

The density tree algorithm [6, 22] of the level set clustering method can not only extract and separate high density clusters of the complex data, but also map the high density clusters into a hierarchical tree structure. The metric of using the “height” at which two points or two clusters merge on the density function is adapted to construct hierarchical tree by density tree algorithm (Figure S1(b)).

### DensityPath Algorithm

DensityPath is developed to reconstruct cell state-transition path of cellular development based on the input of single cell expression profile from scRNAseq. Relying on the density tree of level set clustering method [6, 22], DensityPath analyzes the heterogeneous single cell multimodal density landscape of gene expression space and extracts the underlying cell states of the heterogeneous populations by identifying the high density clusters of RCSs. DensityPath then finds the cell state-transition path by calculating MST of RCSs on the surface of the density landscape. The overview of DensityPath is shown in Figure 1. The main steps for DensityPath are as follows:

A1 **Reduce the dimensionality of scRNAseq data.** DensityPath algorithm first conducts PCA on the data to map data in original space onto the top principal components which preserve the most variance. The choice of dimension embedded in PCA is by cutting off at a component at the inflection point of ordered eigenvalues of PCA, that is the first *i*-th largest eigenvalue *λ*_*i*_ of PCA satisfying

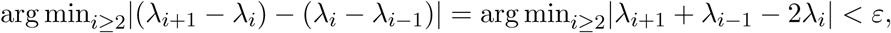

at a given threshold *ε* (e.g. *ε* = 10^−3^ in this study). Following the dimension reduction through PCA, DensityPath then applies the EE algorithm [21] to the embedded PCA data constituted by the top *i*-th largest components onto the *d*-dimensional latent space. We show that *d* = 2 is sufficient to preserve the intrinsic structures of scRNAseq data we studied, even for the data with complex trajectory with branches both from bifurcations and trifurcations. We denote the 2 coordinates from EE algorithm as “EE1” and “EE2” throughout this study.
A2 **Estimate the density function (landscape) and calculate the level sets.** DensityPath estimates the density function (landscape) of the reduced-dimension (low-dimensional) space of single cell expression profile using the standard nonparametric kernel density estimator (KDE) as follows:

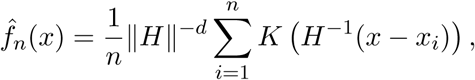

where *H* is the bandwidth and *K* is the kernel. Here the kernel function *K* is chosen as the Gaussian kernel [32]; the choice of bandwidth *H* used in KDE method is calculated based on plug-in method [33, 34]. For any *t >* 0, the estimated *t*-upper level set is calculated as 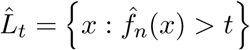. To identify the connected components of 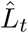, DensityPath first constructs a *k*-NN graph of {*x*_1_, *…, x*_*n*_} on the reduced-dimension space, and then finds the connected components of the subgraph with the nodes restricting to the index set *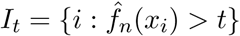*.
A3 **Select high density clusters of RCSs using level set clustering.** DensityPath applies the density tree algorithm of level set clustering method [6, 22] to the density landscape estimated to extract high density clusters as RCSs: by varying threshold *t* and calculating 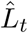 and 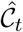, density tree analyzes the structure of the density function to construct a hierarchical structured density tree in an unsupervised manner (see Figure S1(b)). DensityPath takes the high density clusters, which are represented by the external branches of the constructed density tree shown in Figure S1(b), as the RCSs.
A4 **Construct the cell state-transition path.** DensityPath uses the peak points of RCSs as the landmarks, calculates the shortest path (geodesics) of the peak points on the surface of the density landscape. To achieve this, DensityPath applies the Dijkstra’s algorithm to the meshgrid of the density surface as follows: (1) the density surface is represented by the *z*-coordinates (density values) of points above a 2-d square grid in the *x*-*y* plane with uniformly spaced EE1-coordinates and EE2-coordinates; (2) a king’s graph is constructed with the nodes of the grid points, and edges connecting by points with their Moore neighborhood (eight orthogonal and diagonal nearest neighbors) on the 2-d square grid; (3) the distance between each pair of the points is set as (i) the reciprocal of the mean of the two points’ density values if they are connected on the king’s graph or (ii) +*∞* otherwise; (4) given the coordinates of two points (RCSs) on the density surface, their shortest path (geodesics) is calculated by the Dijkstra’s algorithm on the king’s graph. The distortion caused by the difference of distance between the orthogonal connection and diagonal connection on the king’s graph is corrected. DensityPath implements this parts mainly based on the R Package “gdistance” [35]. The embedding cell state-transition path (cell developmental trajectory) is then constructed by finding the MST of the peak points of RCSs.
A5 **Map the single cells onto the cell state-transition path.** To map a given single cell data point onto the cell state-transition path, DensityPath finds a nearest point on the path, which has the smallest geodesic on the surface of the density landscape, to the given cell.
A6 **Calculate the pseudotime of each cell.** If a start cell (root) is determined, the pseudotime of each cell can be calculated as the geodesic distance between the points mapping onto the path of each cell and the initial cell. DensityPath has two tuning parameters. (1) The regularization parameter lambda *λ* in Equation of the EE algorithm, which trades off two terms, one preserving local distance and the other preserving global distance or separate latent points. We show that *λ* = 10 is where the best embeddings occur, and DensityPath is robust when *λ* is in the region from 2 to 50. (2) The number of neighbors *k* for the *k*-NN graphs. The choice of *k* generally relies on the sample size *n*, especially when *n* is small. Here we propose an empirical formulation to compute *k* as *k* = round(*n/*10), that is the nearest integer value of *n/*10. The code of DensityPath is available from https://github.com/ucasdp/DensityPath.

### Method evaluation

To evaluate the pseudotime calculation and branch assignment performance, we adopt two indexes, the Pearson’s correlation coefficient (PCC) and the adjusted rand index (ARI) [36], to the results obtained by different methods on datasets with known branch assignment or real time information of cells. The PCC of the pseudotime calculated and real time information (either experimental time or simulation time) is used to evaluate accuracy or robustness of pseudotime. The ARI, which is a measure of the similarity between two data clustering and has been utilized by Monocle2 [15], is used to measure the accuracy or robustness of trajectory branch assignment of cells. Given a dataset of *n* single cells, and two assignments 𝓧= {*X*_1_, *X*_2_, *X*_*r*_} and *Y* = {*Y*_1_, *Y*_2_, *Y*_*s*_} of the *n* cells into *r* and *s* branches respectively, the overlap between 𝓧 and 𝓨 can be characterized of a set counts *n*_*ij*_ which represents the overlap number in *X*_*i*_ and *Y*_*j*_, i.e. *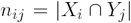*. And we define the number of cells within segment *i* from the former clustering result as *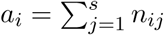*, and the number of cells within the segment *j* from the latter is *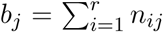*. The ARI value is then formulated as

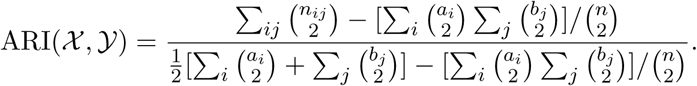

The value of ARI is in the region of [0,1], with a larger value indicating better assignment. In our analysis, the clusters in ARI is defined as the trajectory segment from one branching or initial peaks to the next branching peak point or the end peak.

### Data

We performed DensityPath analysis on three real scRNAseq datasets. (1) The scRNAseq dataset of mouse bone marrow cells is from Paul et al [27]. The processed read count profile of 2730 cells and 8716 genes was provided by authors of [11] (personal communication) and the data was processed into the reads per kilobase per million mapped reads (RPKM) values. We selected out the 3461 informative genes which were provided by authors of [11]. (2) The scRNAseq dataset of mouse hematopoietic stem and progenitor cells (HSPCs) bifurcating to myeloid and erythroid precursors is also from Paul et al [27]. We directly downloaded the processed expression profile of this data by Wishbone [10] which contains 4423 single cells with 2312 informative genes at https://github.com/ManuSetty/wishbone, and we followed the same normalization procedures as Wishbone using their code [10]. (3) The scRNAseq dataset of human preimplantation embryos is from [28]. The expression profile of 1529 single cells with 26178 genes in RPKM values was downloaded from in https://www.ebi.ac.uk/arrayexpress/experiments/E-MTAB-3929/.

We also performed DensityPath analysis on two simulated scRNAseq datasets. The PHATE dataset of the multi-branching trajectories was simulated based on the codes in https://github.com/KrishnaswamyLab/PHATE provided by [20]. The SLS3279 dataset of the bifurcating trajectory was discussed in the supplementary of [29] and the dataset utilized here was from the supplementary material of [37].

## Results

### DensityPath revealed the cluster and developmental structures of bone marrow cells

We tested DensityPath on the scRNAseq dataset of bone marrow cells from [27] (denoted as Paul data), which has combined index fluorescence-assisted cell sorting and been widely used for the study of the heterogeneity in myeloid progenitors. The processed data of the sorted c-Kit^+^Scal^-^lineage(Lin^-^) bone marrow cells, which contains 2730 single cells with 3461 informative genes, has been adopted. In their study, Paul et al [27] identified 19 distinct progenitor clusters at different degree of differentiation through the EM-based clustering approach: accordingly, cluster 1 to 6 represent the erythroid (Ery) fate, cluster 7 to 10 represent the common myeloid progenitor (CMP), cluster 11 corresponds to the dendritic cell fate (DC), and cluster 12 to 18 reflect granulocyte/macrophage progenitor (GMP) fate. Cluster 19, which is constituted by the mis-sorted population of lymphoid progenitors, is denoted as “Outlier” in this study.

DensityPath first mapped the scRNAseq expression profile of 3461 informative genes onto 28 principal components by PCA, and then further mapped the data onto the 2-dimensional subspace of EE1 and EE2 by the nonlinear dimension reduction algorithm EE using the default parameter *λ* = 10 (Figure 2(a))). It is clear that the CMP and their direct progenies, GMP and Ery, are almost unambiguously distinguished into three regions on the 2-d space of EE1 and EE2 by DensityPath, while the DC cells are deviated away from the main trajectory, indicating DC is less relative to the CMP differentiation progression (Figure 2(e))). In addition, DensityPath separated the Outlier cells in cluster 19, a mis-sorted population of lymphoid progenitors, from the progenitor cells of non-lymphoid white blood cells, red blood cells and platelets, exhibiting the validness of our dimension reduction technique (Figure 2(e))).

**Figure 2:**
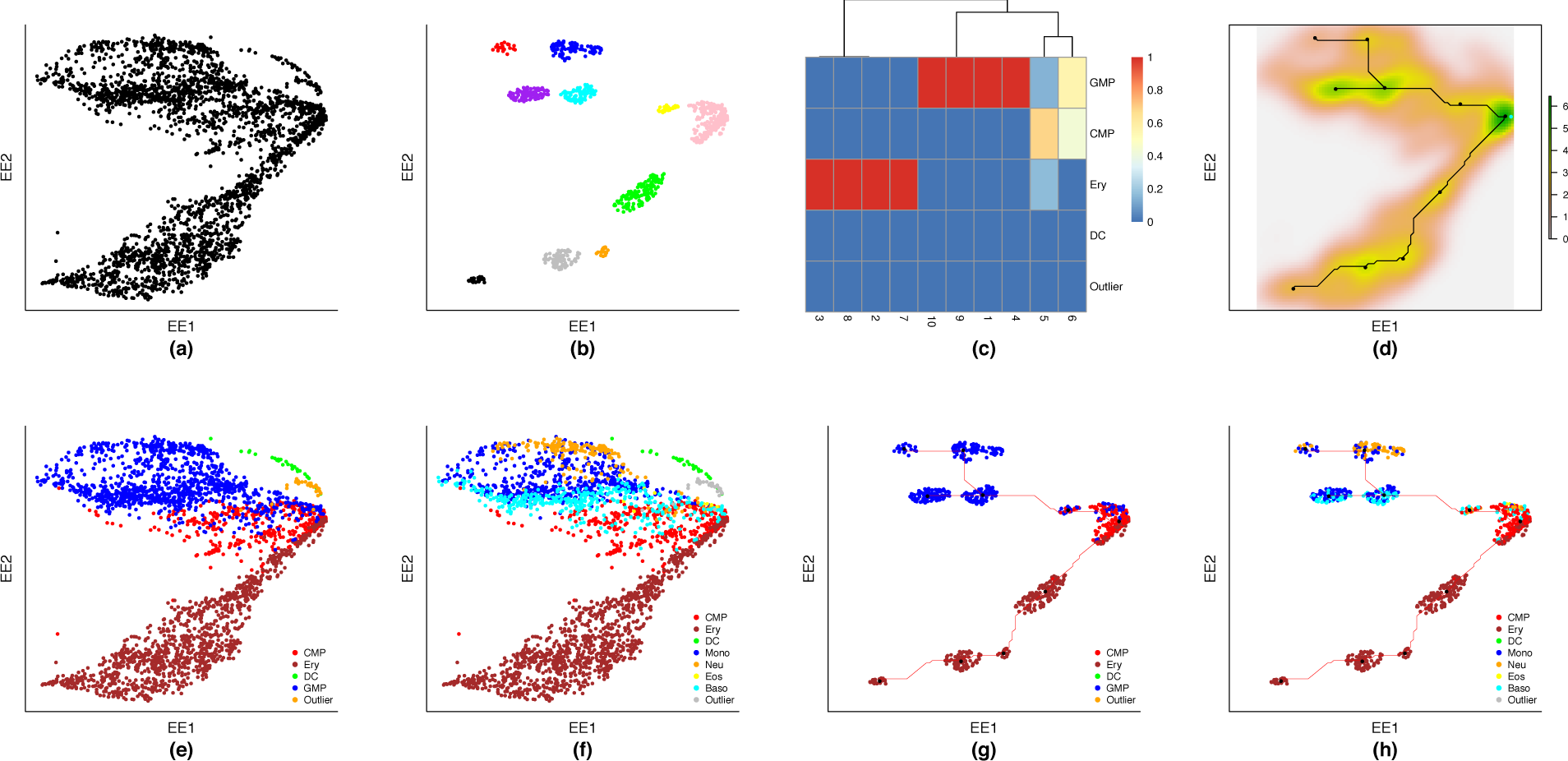
DensityPath revealed the developmental structure of the bone marrow cells.

DensityPath estimated the density landscape of the 2-d of transcriptional heterogeneity of cells by KDE and then conducted the level set clustering analysis, resulting in 10 separate high density clusters of RCSs recognized (Figure 2(b))). The 10 RCSs picked up a total of 865 cells out of the 2730, with the sizes of each cluster ranging from 16 to 276 cells. To reveal the structure of the cellular developmental process, a MST connecting the peak points of RSCs in level set clusters was constructed on the surface of density landscape, showing a cell state-transition path constituted by two bifurcating events, with the first bifurcating event occurring at the right most RCS where the start cell (root) is mapped (Figure 2(d)), the start cell, the first cell of the data, is selected and plotted as the cyan point).

It is worth to noting that, contrasting to the previously defined clusters by Paul et al [27] which is a partition of all cells, DensityPath picked up the high density clusters, and constructed and Outlier by Paul et al [27]. the state transition path connecting density peaks of density clusters. We extracted the 10 high density clusters (RCSs) identified by DensityPath, which show more homologous representation as characterized by the cell clusters defined in Paul et al [27] (Figure 2(g))): among the extracted 10 RCSs, 4 RCSs (Nos. 1,4, 9 and 10) are completely in the GMP clusters; and 4 RCSs (Nos. 2, 3 (99.3%), 7 and 8) are (almost) completely contained by Ery lineage (Figure 2(c))). For the remaining 2 RCSs, the right-most RCS (No. 5) contains cells mainly from CMP (68%) and small portions from Ery (17%) and GMP (15%), showing the occurrence of the bifurcating event that the transition states from CMP to GMP lineage as well as to Ery lineage; the RCS (No. 6), which is the nearest to the right-most RCS (No. 5) from upper-left, contains cells evenly from CMP (44%) and GMP (56%), indicating being in a progression of the transition state from CMP to GMP (Figure 2(c,g)). Due to the scarce number of cells identified as the dendritic cells (30 cells) and the Outlier (31 cells) by Paul et al [27], there are no cells in the RCSs corresponding to the DC and only one cell in the No. 5 RCS corresponding to Outlier, showing that only representative cells in high density clusters are extracted by DensityPath.

DensityPath projected bone marrow cells data in 3461 dimensional space of genes onto 2-dimension plane through PCA and EE techniques. (b) The 10 RCSs were extracted by level set clustering: the indexes for the RCSs followed by their corresponding colors and cell numbers are 1 (red, 22), 2 (black, 26), 3 (green,130), 4 (blue, 96), 5 (pink, 276), 6 (yellow, 25), 7 (grey, 80), 8 (orange, 16), 9 (purple, 104), and 10 (cyan, 90). (c) The heatmap shows the percentages of cells in each of 10 RCSs (x-axis) that are distributed into the clusters CMP, Ery, DC, GMP and Outlier by Paul et al [27] (y-axis) respectively. (d) A branching trajectory was obtained by constructing the MST of density peaks in RCSs using the geodesic of density surface. Given the start cell marked in the plot with cyan, this is a progression with two bifurcating events. (e,g) All cells (e) and the cells in the 10 RCSs (g) are plotted on the EE1 and EE2 coordinates, showing a separation of clusters CMP, Ery, DC, GMP and Outlier by Paul et al [27] with progression trends. (f,h) are the same as (e,g), except using detailed clusters information as CMP, Ery, DC, Mono, Neu, Eos, Baso, Outlier by Paul et al [27].

Noting that DensityPath constructed 2 branches on the trajectory of the GMP lineage (upper-left of Figure 2(g))), showing a second bifurcating event. We thus further divided the GMP cluster into 4 cellular subtypes/lineages as monocytes (Mono), neutrophils (Neu), eosinophils (Eos), and basophils (Baso) according to Paul et al [27]: the 4 subtypes showed clear transition patterns of developmental progression (Figure 2(f)) and Figure S2). When looking at the cells of the RCSs along the GMP lineage, the 4 sub lineages maintain their continuity and the progression order of cell types: Neu and Baso fate are clearly separated in RCSs from the two branches of the second bifurcating events, while Mono can be found on both branches (Figure 2(h))).

### DensityPath reconstructed the branched trajectories of HSPCs bifurcating to myeloid and erythroid precursors

We applied DensityPath to another scRNAseq dataset generated by [27], consisting of single cells extracted from the process of mouse HSPCs bifurcating to myeloid and erythroid precursors (denoted as HSPCs data). Wishbone algorithm worked on the HSPCs data to reconstruct the bifurcating developmental trajectories [10]. We directly downloaded the preprocessed expression profile of HSPCs data in [10] which contains 4423 single cells with 2312 informative genes, and normalized the data as [10].

DensityPath first mapped the scRNAseq expression profile of 2312 informative genes onto 6 principal components by PCA, and then further mapped the data onto the 2-dimensional subspace of EE1 and EE2 by EE using the default parameter *λ* = 10 (Figure 3(c))). DensityPath estimated the 2-d density landscape of single cells by KDE approach, and then conducted the level set clustering analysis, which result in 9 separate high density clusters of RCSs, with sizes ranges from 49 to 660 that summing up to 2178 of the total 4423 cells (Figure 3(a))). To reveal the structure of the cellular developmental process, DensityPath connected the peak points of RSCs by the MST on the density landscape surface, as the cell state-transition path. It is clear that the 9 RCSs are aligned along a branched-shape trajectory which coincides with the actual bifurcating process form HSPCs to myeloid and erythroid precursor as well as the result out of Wishbone algorithm [10] (Figure 3(b))).

**Figure 3:**
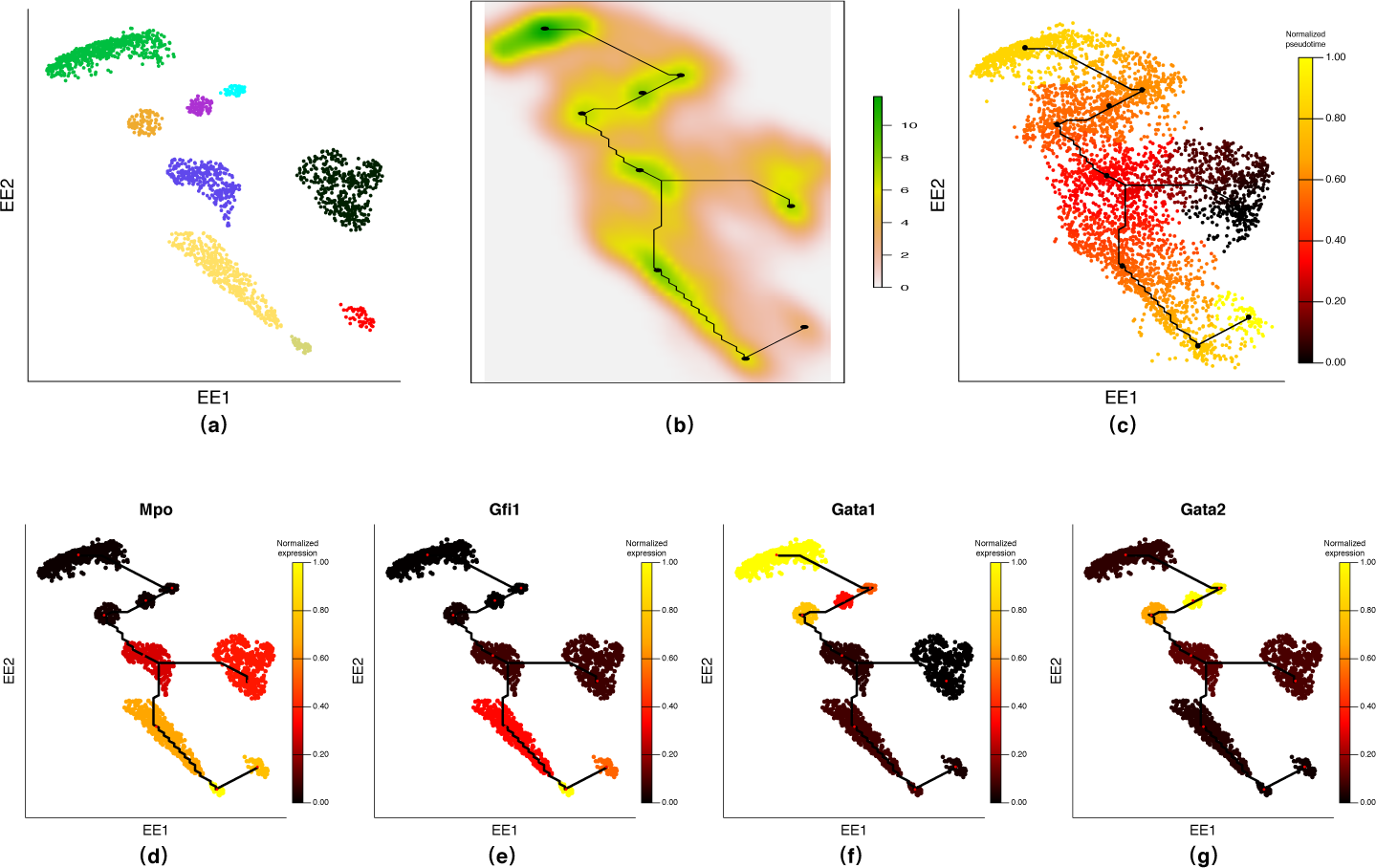
DensityPath recovered the bifurcating structure of HSPCs data. (a) DensityPath extracted 9 separate RCSs and cells of each RSC have different colors. (b) A bifurcating trajectory was constructed by connecting the peaks of the RCSs. (c) Given the same start cell (the 1432-nd cell of the data) as Wishbone, DensityPath calculated the pseudotime of each cell, with time of the start cell setting as 0. (d)-(g) The expression of the 4 marker genes (*Mpo, Gfi1, Gata1* and *Gata2*) of myeloid or erythroid lineage are plotted on the cells of the RCSs along the trajectory. The mean value of expression of each gene in the cells from each RCSs were calculated and shown on the cells of the corresponding RCSs. The two branches show opposite expression pattern on the 4 marker genes *Mpo, Gfi1, Gata1* and *Gata2*.

To calculate the pseudotime, we first selected the same start cell as in [10]. We then calculated the geodesic between start cell and the other cells, all of which have been mapped onto the MST path, on the density landscape as our pseudotime: the black points on the middle-right regions is identified as trunk, while points with colors ranging from slant-red to yellow on the up-left and down-left regions reflect two branches with opposite directions (Figure 3(c))). In addition to the similar bifurcating structure, the points in density clusters that distributed on the trunk and branches are mostly in accordance with the result of Wishbone [10]. In detail, after mapping all cells onto the cell state-transition path, there are 952 cells, which account for 98.65% of 965 cells in the trunk of DensityPath result, located in the trunk defined in Wishbone; 1253 cells in the branch 1 obtained by Wishbone, taking 87.81% share of 1427 cells in one of the main branches of DensityPath; and 1915 cells in the branch 2 of Wishbone, accounting for 94.29% of 2031 cells located in the other main branch of DensityPath. The PCC for the pseudotime calculated by DensityPath with geodesic described above and in Wishbone algorithm is 0.83.

The expression trends of the marker genes for the myeloid lineage and the erythroid lineage are shown along the reconstructed trajectory (Figure 3(d,e,f,g)) and Fiurge S3): The myeloid markers of *Mpo, Gfi1* and *Cebpe* as well as the erythroid markers of *Klf1, Gfi1b, Gata1* and *Gata2*, were selected and shown distinct trend along the two branches, indicating that the upper-left branch corresponds to the erythroid lineage while the other one corresponds to the myeloid lineage. Furthermore, *Gata2* showed an earlier activation with high expression along the erythroid lineage than that for *Gata1* (Figure 3(f,g)), consisting with our current understanding [10, 38]

### DensityPath reconstructed the complex trajectory with both bifurcating and trifurcating events on human preimplantation embryos

We applied DensityPath to another scRNAseq dataset of 1529 single cells of human preimplantation embryos from Petropoulos et al [28] (denoted as HPE data). The cells isolated from embryos ranging from the 8-cell stage up to the time-point just prior to implantation were collected and labeled with time from embryonic day 3 to day 7 (E3 to E7). The differentiation process of the cells of human preimplantation embryos was characterized as the synchronous differentiation of the inner cell mass (ICM) into three distinct cell types of the mature blastocyst: trophectoderm (TE), primitive endoderm (PE) and epiblast (EPI) at E5 from the zygote [28]. Besides the dominant processes of the segregation of ICM, the cells in TE lineage were also reported to be further subdivided into two subpopulations, reflecting that polar and mural cells are present within the TE lineage [28].

We filtered less variable genes out and selected the 4600 most variable ones across all cells as in [14] for further DensityPath analysis. PCA and EE were successively applied with default settings to map the expression profile in high-dimension space onto 2-d plane of EE1 and EE2. DensityPath then estimated the density landscape by KDE method, and picked up the 14 high density clusters of RCSs with sizes ranges from 1 to 84, summing up to 500 of the total 1529 cells. Finally, DensityPath connected the peak points of RSCs by the MST on the density landscape surface, resulting in a complex trajectory with 2 bifurcating events and 1 trifurcating event (Figure 4(a))). The trifurcating event occurring at E5 identified by DensityPath is completely consistent with the progression reported in [28], that ICM lineage separates into PE and EPI lineages following the segregation of zygote into TE and ICM lineage at same day of E5. The bifurcating event appeared at E6 and E7 is also consistent with [28] that two subpopulations occurred in TE lineage. The bifurcating event occurring at E4 is still not clear, deserving further study. The PCC for the pseudotime calculated by DensityPath with the experimental embryonic time of the cells is 0.8153.

**Figure 4:**
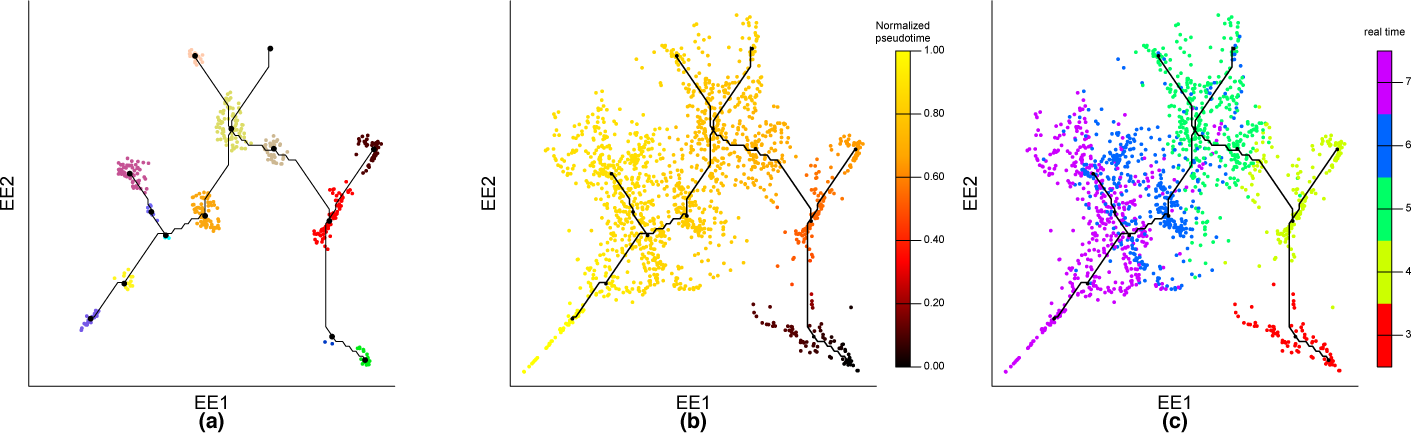
DensityPath reconstructed the complex trajectory of the HPE data. (a) DensityPath reconstructed a complex trajectory with 2 bifurcating events and 1 trifurcating event by connecting the peaks of the 14 separate RCSs identified by level set clustering. (b) Given start cell (the 1472-nd cell of the data), DensityPath calculated the pseudotime of each cell, with time of the start cell setting as 0. (c) The embryonic day (E3-E7) of the cells are plotted different colors on the cells ordering along the trajectory by DensityPath.

### Validation of DensityPath on simulated datasets

We tested DensityPath on two simulated datasets with varying numbers of branches on the trajectories and varying sample sizes to assess the accuracies in the dimension reduction, pseudotime calculation as well as the branch assignment.

We first tested DensityPath on the simulated tree dataset with a total of 1440 single cells and 60 genes from [20] (denote as PHATE data): the simulated tree has 1 trunk and 9 branches, each of which contains around 100 points (cells) in different data subdimensions of a 60 dimensional space to model a system where the progression along a branch corresponds to an increasing expression of a couple of genes. The tree has three bifurcating events and one trifurcating event: the trunk (red) first bifurcates into two branches (green and black), one (green) of which subsequently bifurcates into two branches, while the other one (black) trifurcates into three branches, and following by one (blue) of the three branches further bifurcating into two branches (see the ground truth of the data in Figure 5(a)), which is based on Figure 7(a) of [20]). DensityPath mapped the data onto the 2-d plane of EE1 and EE2, and reconstructed the complex trajectory with 1 trunk and 9 branches with the same tree structure, recovering the simulated progression fully (Figure 5(b))). With the initial cell fixed as the start point of simulated time, we calculate the pseudotime of the cells (Figure 5(c))). The PCC of the pseudotime recovered by DensityPath and the real time of the simulation is 0.9528, while the ARI value between the branch assignment of the cells to the reconstructed trajectory by DensityPath and the ground truth by simulation is 0.7317.

**Figure 5:**
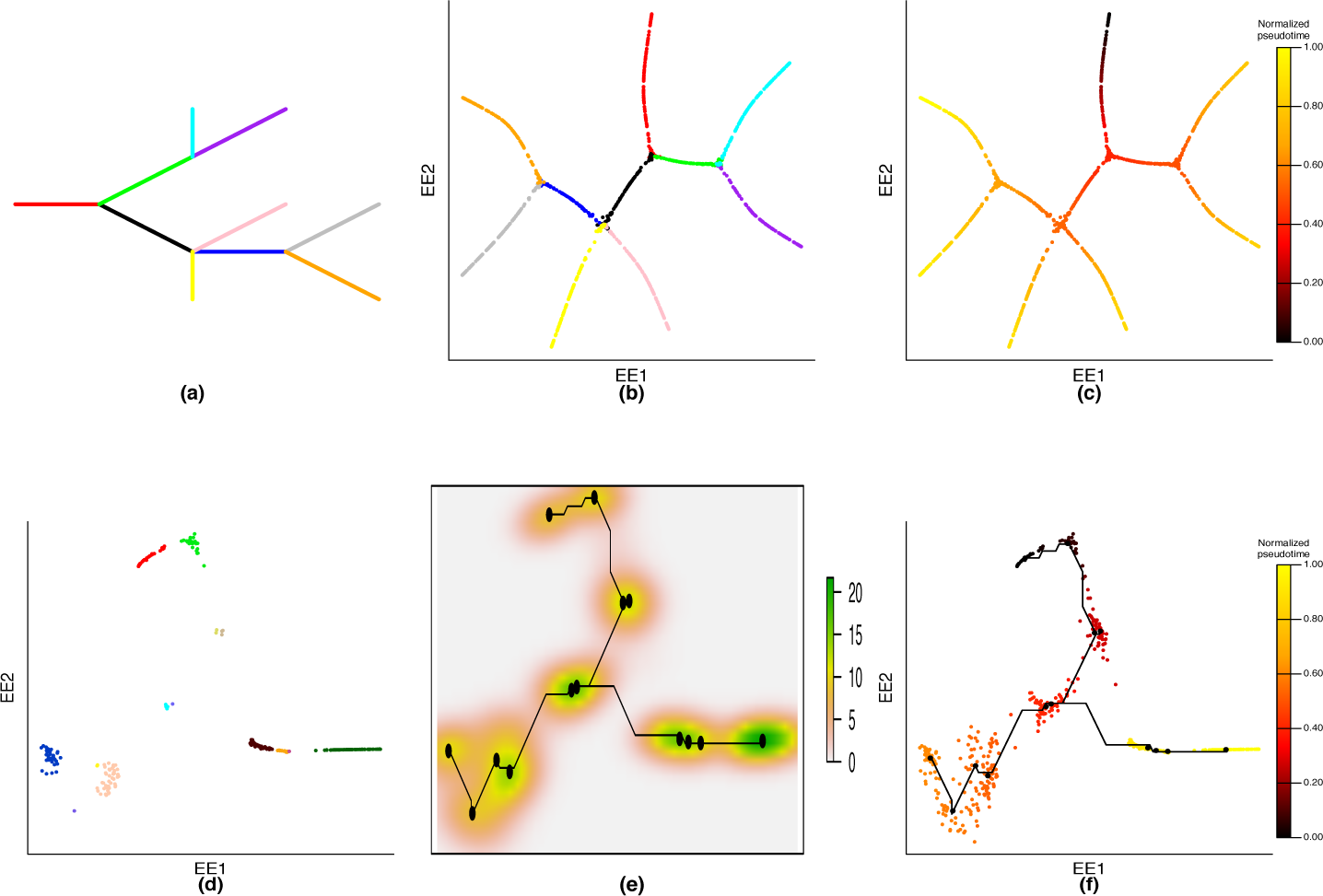
Validation of DensityPath on the simulated PHATE data and SLS3279 data. (a) The ground truth of PHATE data based on Figure 7(a) of [20]). (b) DensityPath mapped the PHATE data onto the 2-d plane of EE1 and EE2, and reconstructed the complex trajectory, accurately recovering the structure of PHATE data. (c) DensityPath calculated the pseudotime of the PHATE data, by fixing the initial cell in simulation (the 1-st cell of the data) as start cell. (d) DensityPath mapped the SLS3279 data onto the 2-d plane of EE1 and EE2, and extracted 14 RCSs. (e) DensityPath reconstructed the cell development bifurcating trajectory on density landscape of the SLS3279 data. (f) DensityPath calculated the pseudotime of the SLS3279 data, by fixing the initial cell in simulation (the 1-st cell of the data) as start cell.

We further tested DensityPath on the simulated dataset with 490 single cells and 48 genes from [29] (denoted as SLS3279 data), which models a bifurcating trajectory data with two terminating destinations. DensityPath recovered a branching structure on the 2-d space of EE1 and EE2 by PCA and EE, and extracted a total of 287 cells into 14 RCSs (Figure 5(d,e)). We also fixed the initial cell (Cell 1) which was selected according to the simulated time, as the start cell and calculated the pseudotime by DensityPath (Figure 5(f))). The PCC of the pseudotime calculated by DensityPath and the time in simulated progression is 0.9291. The result indicates that even with limited sample size (*<* 500), which is unfavorable to KDE, DensityPath can still accurately recover the tree trajectory.

## Comparison with other methods

### Paul data

We compared DensityPath with Monocle2, DPT, Wishbone, and TSCAN on our processed Paul data [27]. (1) Monocle2 first excluded the mis-sorted cells (Outlier) based on the prior information of the cells, identified 12 states, and revealed a branched structure with two bifurcating events (Figure S4(a)). Although the GMP and Ery cells were correctly separated into 2 branches by Monocle2, a nonnegligible number of CMP cells are assigned together along the branch of GMP (Figure S4(b,c)). (2) DPT method was applied 3 times consecutively to the Paul data to identify multi bifurcating events, resulting in a trajectory with 7 branches identified (Figure S5(a)). The 7 branches cannot fully reveal the CMP, Ery and GMP lineages, and the Outlier cells were assigned to the branch2, mixing with cells in CMP and Ery (Figure S5(a)). (3) Wishbone reconstructed a bifurcating structure with one branch point on Paul data: 560 cells is assigned to the trunk, 985 cells is assigned to branch 1, and 1185 cells is assigned branch 2 (Figure S6(a)); About 2/3 CMP cells are located on trunk part while the GMP and Ery cells are located on branch 2 and branch 1 respectively, achieving a high quality of assignment (Figure S6(b)). However, the Outlier cells cannot be identified and are clustered onto the trunk together with DC cells, which should be independent from the progression of CMP development (Figure S6(b,c)). (4) TSCAN generated 8 clusters, with 2 clusters mostly in GMP, 1 cluster completely in Ery and 2 clusters mostly in Ery, 2 clusters mostly in CMP and one overlap mainly in GMP and CMP (Figure S7(a,b)). When considering the detailed cell type clustering result, TSCAN tends to combine the lineages in GMP closely while the mis-sorted cells are also distributed along the main trajectory, failing to distinguish the outliers from the progression of non-lymphoid cells.

### HPE data

We applied the Monocle2, DPT, and Wishbone to the HPE data [28] for the comparison of the pseudotime calculation with the known experimental embryonic time of cells. For the calculation of pseudotime, the same start cell as for DensityPath was assigned to all methods. Since Monocle2 algorithm failed in results for this dataset, we only compared the results obtained in DPT and Wishbone to DensityPath. (1) DPT reconstructed a bifurcating structure where 1399 cells are allocated to branch2, 85 and 29 cells distributed into branch1 and branch3, respectively, with 16 cells undefined (Figure S8). The cells in E5 and E6 are tightly collapsed by DPT, and the trifurcating event as clarified in [28] is hard to be observed by DPT. The diffusion pseudotime of DPT has a PCC of 0.6189 with the experimental embryonic time, which is lower than DensityPath (0.8153). (2) Wishbone revealed a bifurcating structure with 183 cell on trunk, 1289 cells on branch1 and 57 cells on branch2. The branching structure (Figure S9) assigns the cells from E3 and part of from E4 to the trunk part, the others from E4 to branch1 and almost all cells from E5-7 to the branch2 part. With the branching part identified between E4 and E5, the main differentiation process at E5 and E6, E7 are not revealed by Wishbone. The pseudotime obtained in Wishbone has a PCC of 0.8363 with the experimental embryonic time, which is marginally higher than that of DensityPath (0.8153).

### Simulated datasets of PHATE and SLS3279

We also compared DensityPath with Monocle2, DPT and Wishbone on the simulated PHATE data with complex trajectory with both bifurcating and trifurcating events [20], Compared with the ground truth of the data (Figure 5(a))), none of Monocle2, DPT and Wishbone fully reconstructed the branching structure (Figure S10 and Figure S11). We computed the ARI values for each branch assignment obtained, DensityPath outperformed the other methods with ARI value of 0.7317, while none of the others methods obtained ARI greater than 0.5 (Table 1). In addition to contrast the consistency for branch assignment, we also calculated the pseudotime using the comparing methods by fixing the first cell as start point, and calculated PCC in terms of the actual time reported in the simulation data. DensityPath again performed best among the others with the PCC as high as 0.9528 (Table 1).

**Table 1:** The comparison of PCC and ARI between different methods on simulated datasets of PHATE and SLS3279. Since no real branch assignment information is available for the SLS3279 data, no ARI values for the SLS3279 data are calculated (denoted as “NA”).

|  | PHATE |  | SLS3279 |  |
| --- | --- | --- | --- | --- |
| Method | PCC | ARI | PCC | ARI |
| Monocle2 | 0.8127 | 0.4427 | 0.8602 | NA |
| Wishbone | 0.8613 | 0.2286 | 0.8888 | NA |
| DPT | 0.8663 | 0.4076 | 0.9270 | NA |
| DensityPath | 0.9528 | 0.7317 | 0.9291 | NA |

Since DPT and Wishbone are designed and optimized to reconstruct the trajectory with one branch point, we thus compared DensityPath with these methods together with Monocle2 on simulated data SLS3279 in [29]. All the methods recovered the known branched structure (Figure S12, Figure S13, and Figure S14). Besides, Monocle2 also revealed a short tiny branch in one of the end of main branches (Figure S12). When comparing the PCC of pseudotime and simulated time, DensityPath has the highest PCC of 0.9291 and DPT has a second highest PCC of 0.9270 (Table 1). Since no branch assignment information is available for the simulated SLS3279 data, we did not compare the ARI values.

### Robustness analysis of DensityPath

There is a user-defined parameter in DensityPath algorithm, *λ*, which serves as a regularization term when calculating the embedded algorithm EE. It trades off the terms of preserving local distances with global distances or separate latent points. The DensityPath algorithm with *λ* varying from 2 to 50 were tested on the PHATE data. The according values of PCC and ARI, though with small fluctuation, are robust (Figure 6(a))). In addition to the robustness of parameter choice in EE, we tested the robustness of DensityPath on the level set clustering method by resampling subsets of samples of PHATE data after the dimension reduction of A1 step of DensityPath to calculate the pseudotime and branch assignment. With the proportion of resampling scale varying from 0.1 to 0.9, the results of PCC on pseudotime and ARI on branch assignment were quite stable in median values with the decrease of variances (Figure 6(b,c)).

**Figure 6:**
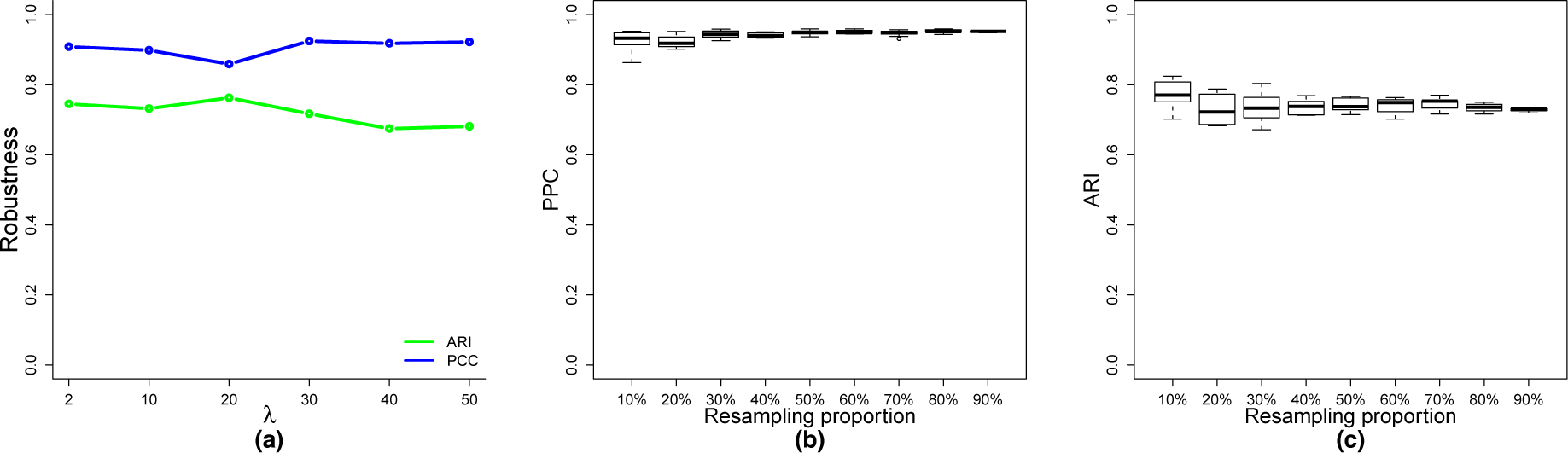
The robustness of DensityPath on the parameter choices and sample resampling. (a) The values of the PCC and ARI of DensityPath by the choices of *λ* on PHATE data. (b) The boxplot of PCC values of DensityPath by resampling different proportions of PHATE samples. (c) The boxplot of ARI values of DensityPath by resampling different proportions of PHATE samples.

## Discussion

In this study, we proposed the DensityPath algorithm that accurately and efficiently visualizes and recovers the unknown cell developmental trajectories for large-scale scRNA-seq data. DensityPath outperformed existing methods such as Monocle2, DPT, Wishbone and Wishbone by adopting EE algorithm for the nonlinear dimension reduction, applying a powerful level set method to characterize the heterogeneous single cell density landscape of gene expression space and utilizing the geodesic on the density landscape to metric cell state transition path.

The dimension reduction plays a vital role in the scRNAseq data analysis. In this study, we showed that EE can preserve the intrinsic structure of both global and local nonlinear manifold hidden in the data, and visualize the complex trajectories in 2-d plane of EE1 and EE2. Apart from EE, other dimension reduction techniques could also be used to scRNAseq, whereas to the best of our knowledge EE is the one that generally suits best for the density based analysis of level set approach in our study. We applied PCA, diffusion map, tSNE, PHATE and EE to HPCSs data [10] and the simulated PHATE data [20], and then performed level set clustering algorithm to construct differentiation trajectory, respectively. The structure identified by the five dimension reduction techniques varies significantly on PHATE data [20] (Figure S15). In view of the HPCSs data [10] the trajectories obtained by diffusion map and PHATE is one-dimensional with no branching points recognized (Figure S15), while in view of the PHATE data the multi-branching structures obtained by PCA, tSNE and diffusion map are quite confused (Figure S16).

Many cell developmental trajectory reconstruction or visualization methods, including TSCAN and Monocle2/RGE, rely on using the clustering methods such as hierarchical clustering and *K*-means algorithm to construct the principal graphs. However, these clustering methods, which seek an optimal partition of the data, may be unstable when the data are heterogeneous and noisy or exhibit complex multimodal structure. Furthermore, the both clustering methods require pre-specification of the number of clusters, and the output maybe sensitive to the number of *K* specified [15]. On the contrary, DensityPath applied the level set clustering method to extract the RCSs, which has superior for the clustering and characterizing the complex heterogeneous single cell data. DensityPath does not require pre-specification of the number of clusters, and is robust to the choice of the tuning parameters.

The choice of the bandwidth *H* of KDE in the DensityPath algorithm plays an important role. Small bandwidths give very rough estimates while large bandwidths give smoother estimates [32]. Although many methods such as rule of thumb, least square cross-validation, biased cross-validation, and plug-in method have been developed on the choice of bandwidth *H*, it remains an open question regarding how to optimally choose the smoothing bandwidth that lead to better estimate of geometric/topological structures [39]. Nontheless, we found that the plug-in method [33, 34] is superior to other methods in our study.

The theoretical framework and justification of the Waddington landscape was solved for a gene network constituted by two interacting genes [25]. However, for gene networks with tens of thousands of genes, it still remains challenge to understand the mechanisms of epigenetic landscape. Recently, the physical model based methods are emerging to construct energy landscape and transition state path using the scRNAseq data [37, 40]. These methods are mainly validated on datasets with sample size less than 1000, and remain unclear about their performances on large-scale datasets. On the other hand, DensityPath does not reply any specific physical model in the dimension reduction step and applied nonparametric method to construct the density landscape, which is a transformation of the the energy function, in a data-driven fashion. DensityPath is efficient to handle large-scale datasets. Further study of the connection of the density landscape and the physical model based energy landscape will be an interesting topic for our future work.

Recently, time series scRNAseq data are accumulating, providing a grand challenge for computational biologists to understand the dynamics of the cellular reprogramming. The pioneer work has applied sophisticated mathematical tool of optimal-transport analysis to model the data [41]. The level set method will also exhibit great potential for the analysis of developmental landscape of time series scRNAseq, and it will also be an interesting topic for our future work.

## Acknowledgements

This work is supported by the Strategic Priority Research Program of the CAS (XDB13050000), the NCMIS of the CAS, the LSC of the CAS, the Youth Innovation Promotion Association of the CAS, and NSFC grants (Nos.11571349, 91630314). LW would like to thank the Mathematical Biosciences Institute (MBI) at Ohio State University, for partially supporting this research. MBI receives its funding through the NSF grant DMS 1440386.

## Appendix Supplementary Figures

**Figure S1:**
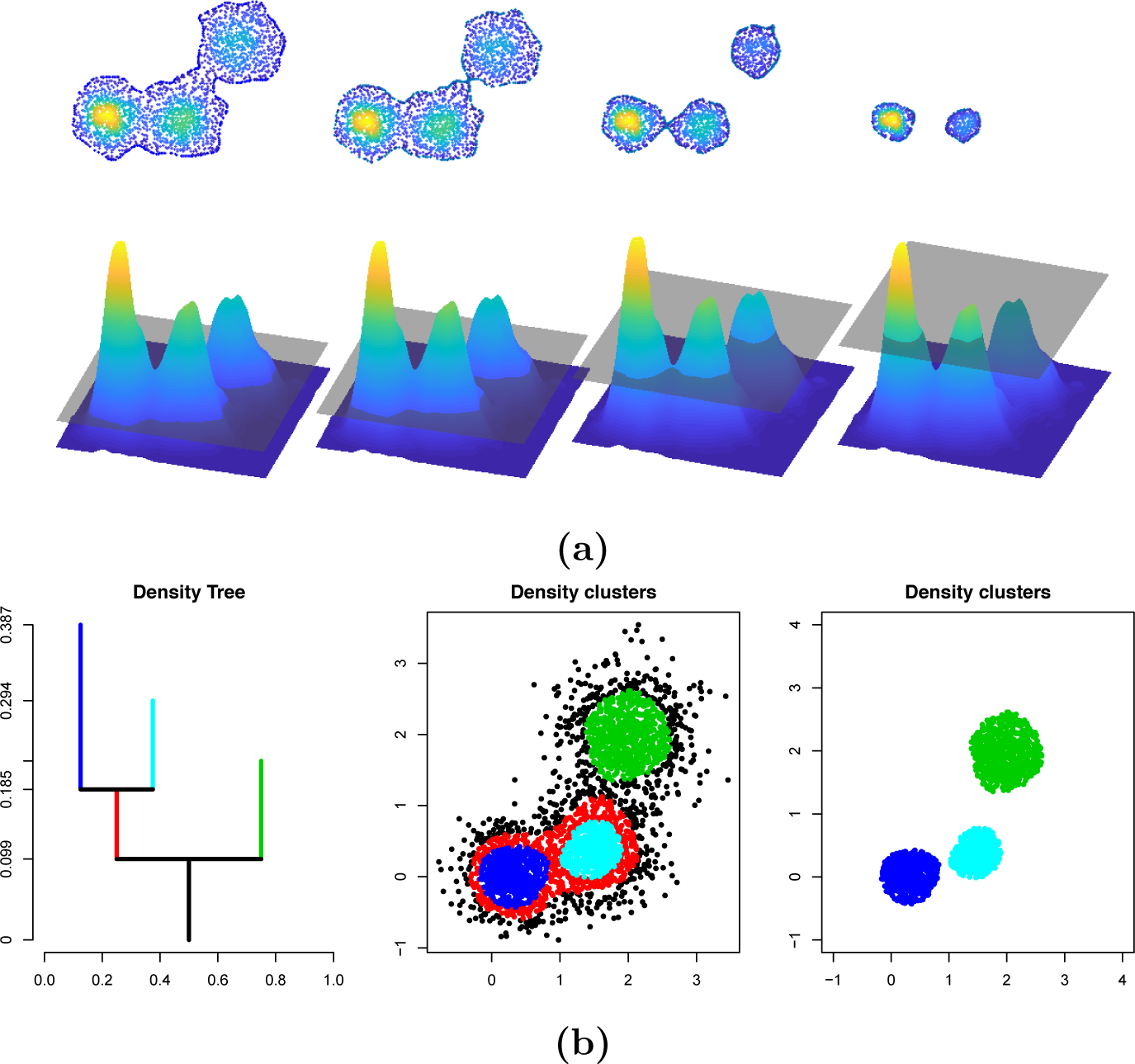
Overview of Level Set Cluster. (a) DensityPath estimates the single cell density landscape (function) of the reduced-dimension space of gene expression (here *d* = 2); as increasing the threshold *t*, DenistyPath analyzes the structure of the density landscape by calculating the *t*-upper level sets 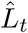 and the connected component(s) 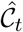. (b) DensityPath applies density tree algorithm to the level set results to construct the density tree (Left), cluster the single cells according to the density tree (Middle) and select the separate high density clusters (Right).

**Figure S2:**
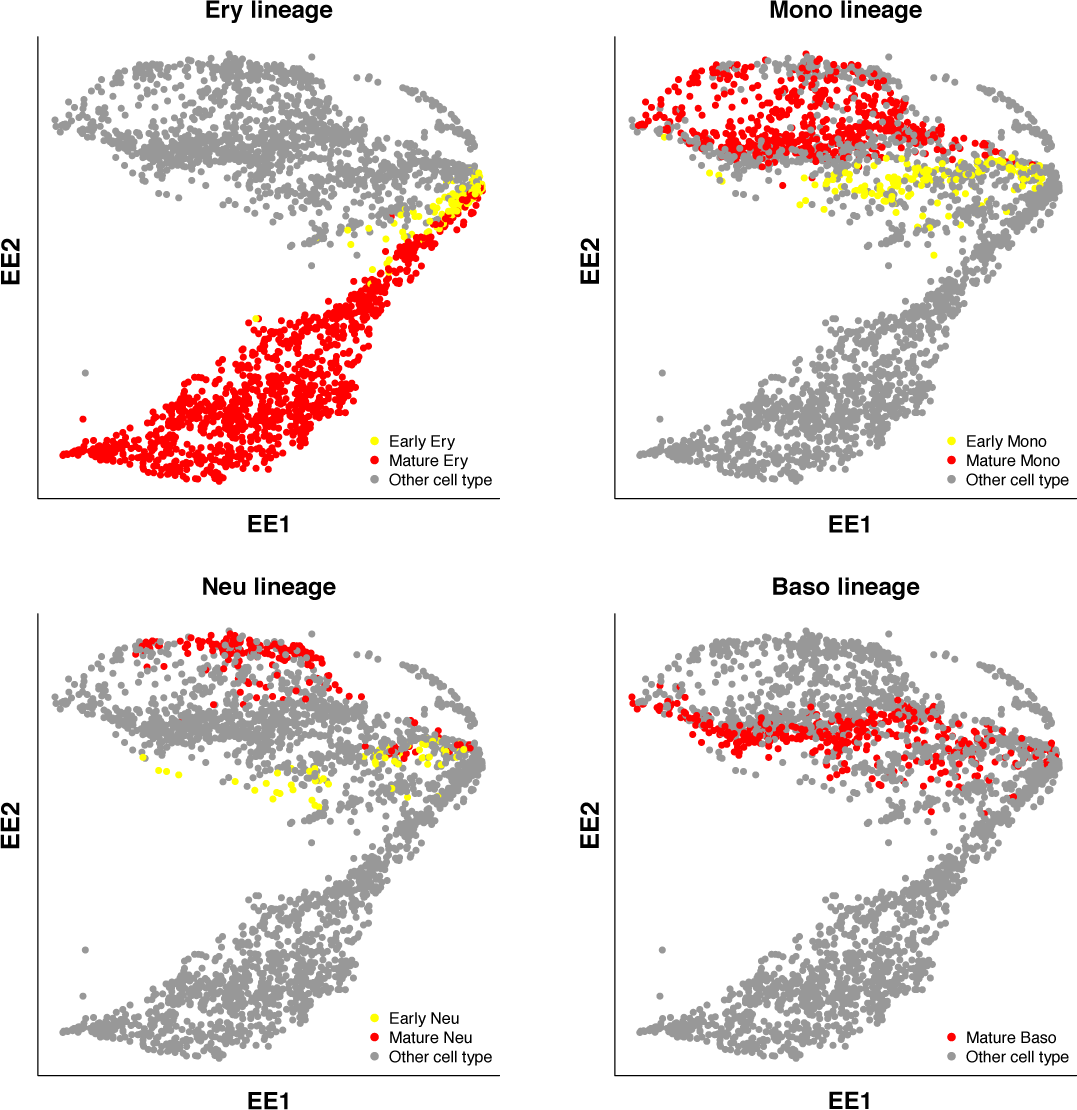
Scatter plots of 4 lineages in Paul data on the 2-d plane of EE1 and EE2.

**Figure S3:**
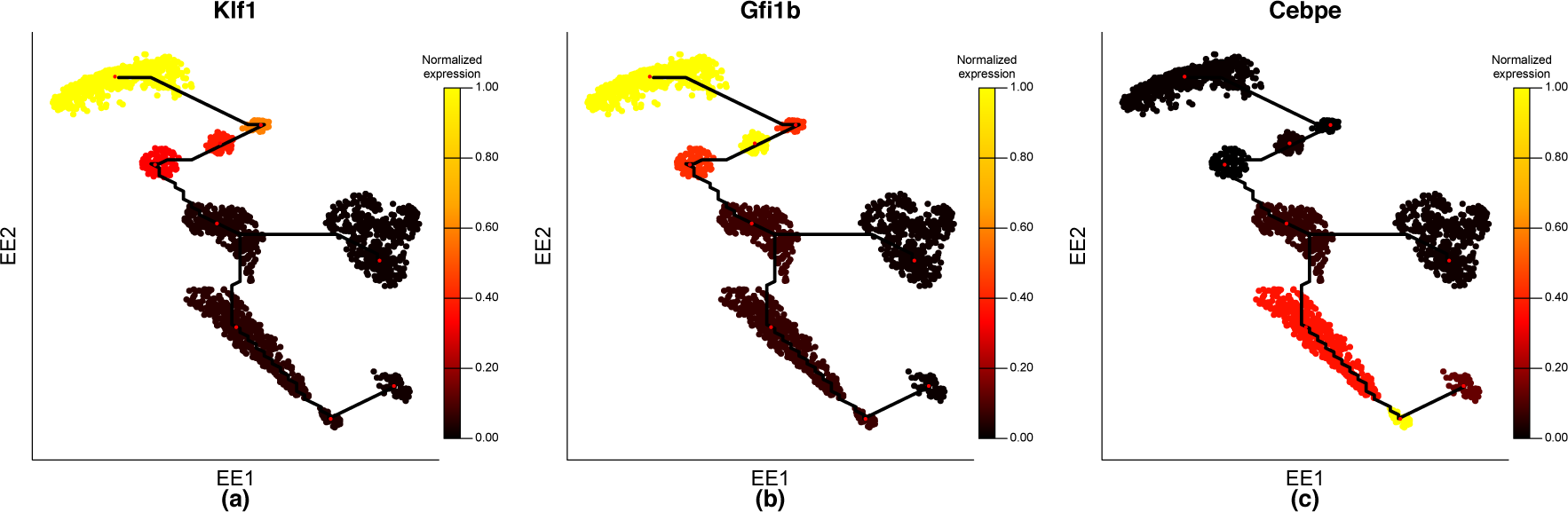
The expression of genes *Klf1 Gfi1b* and *Cebpe* along the trajectory of the HSPCs data. The black curves represent the trajectory reconstructed by DensityPath algorithm. The expression of the 3 marker genes *Klf1 Gfi1b* and *Cebpe* of myeloid or erythroid lineage are plot on the cells of the RCSs along the trajectory. The mean value of expression of each gene in the cells from each RCSs are calculated and shown on the cells of the corresponding RCSs. The two branches show opposite expression pattern on the 3 marker genes.

**Figure S4:**
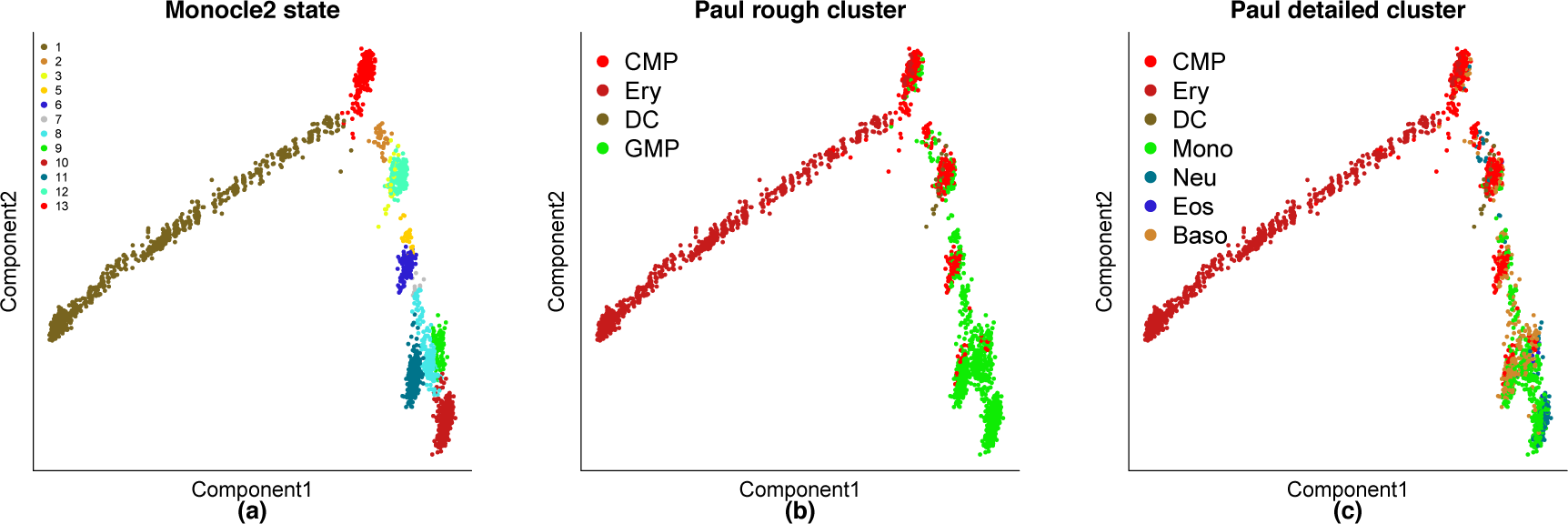
Monocle2 analysis on Paul data. Monocle2’s code “Paul dataset analysis final.html” (from online supplementary script of [15]) was performed on our processed Paul data. (a) Monocle2 removed Outlier cells based on the prior cell information, identified 12 states, and reconstructed a multi-bifurcating trajectory. (b) The distribution of the main four cell types. (c) The distribution of cells with detailed classification on the 2-dimension space.

**Figure S5:**
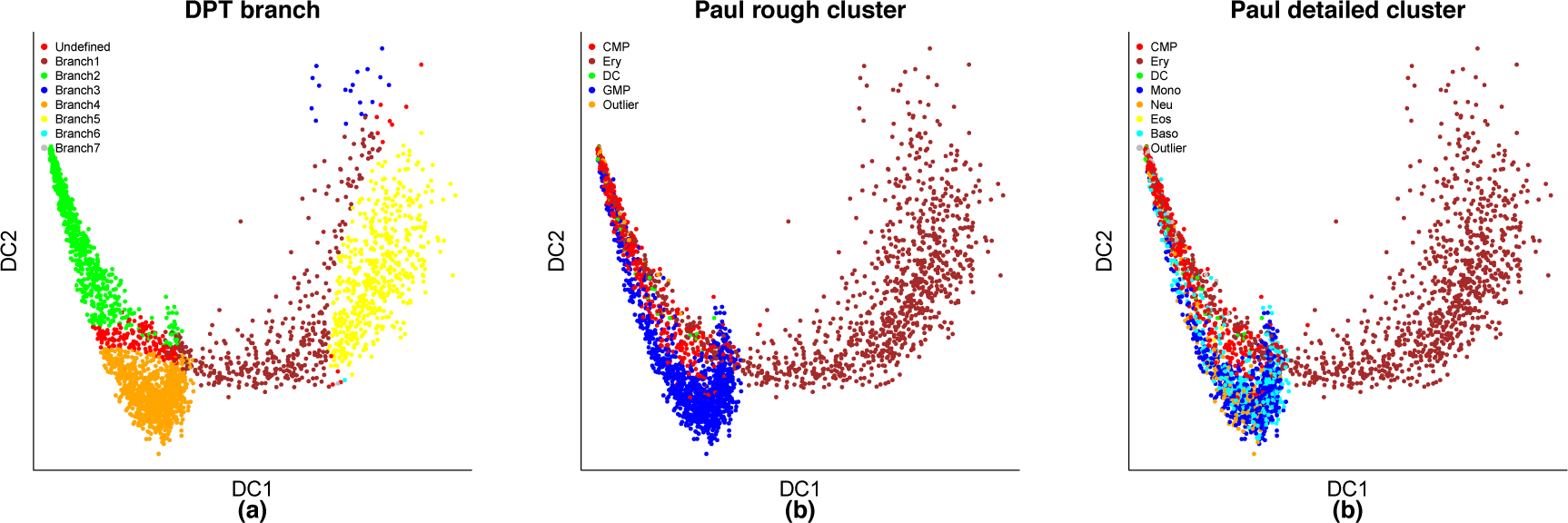
DPT analysis on Paul dataset. The DPT code “run a general example.m” downloaded from supplementary material of [11] was performed by setting the parameter sigma as 20. We carried out 3 times DPT analysis: We first applied DPT to the whole cells and obtained 3 branches with the number of cells 2182, 522 and 18 respectively; then we performed DPT analysis on branch 1 (2182 cells) and branch 2 (522 cells) separately, resulting a total of 7 branches. (a) DPT reconstructed a bifurcating trajectory. The visualization of results in DPT is demonstrated through Diffusion Map. (b) The distribution of the main five cell types. (c) The distribution of cells with detailed classification on the 2-dimension space.

**Figure S6:**
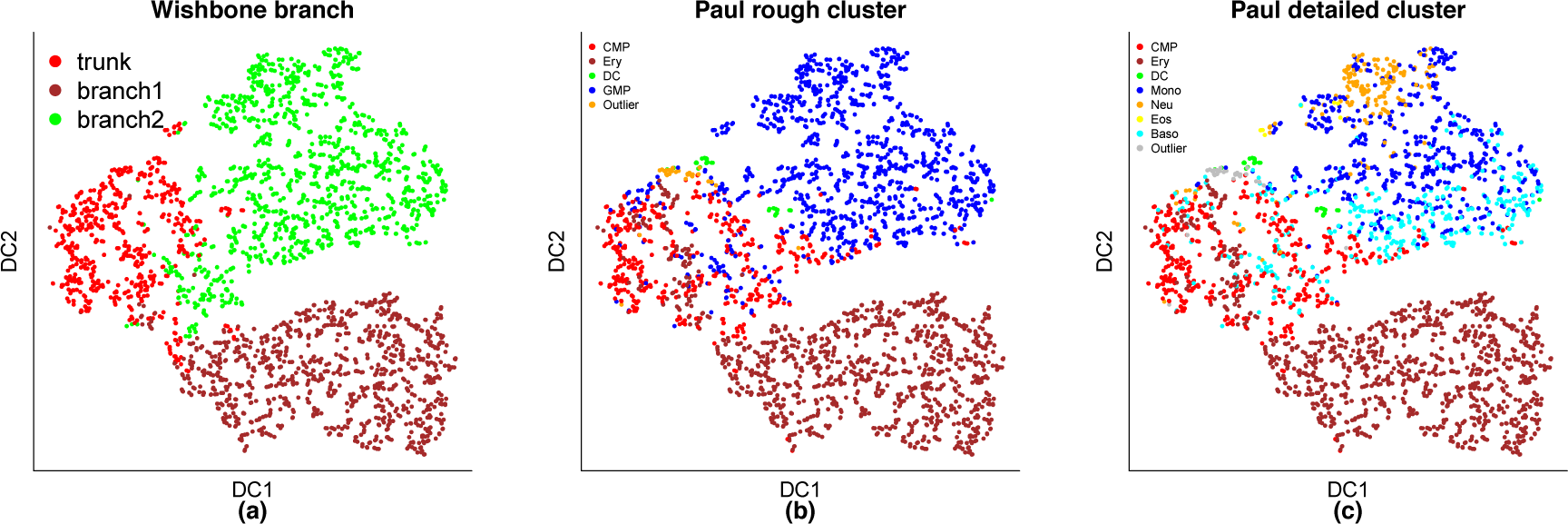
Wishbone analysis on Paul data. The Wishbone was performed by setting the parameters pca and k as 15 and 15, respectively. The visualization of results in Wishbone is demonstrated through tSNE. (a) Wishbone reconstructed a bifurcating trajectory. (b) The distribution of the main five cell types. (c) The distribution of cells with detailed classification on the 2-dimension space.

**Figure S7:**
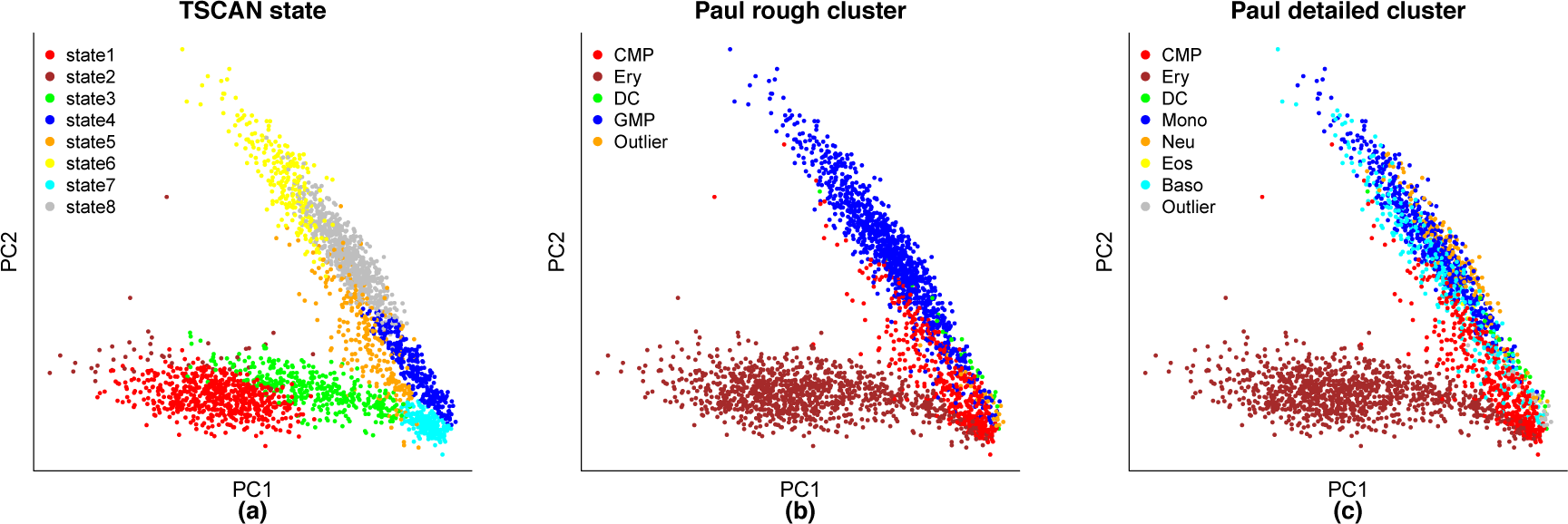
TSCAN analysis on Paul data. (a) TSCAN reconstructed a bifurcating trajectory. (b) The distribution of the main five cell types. (c) The distribution of cells with detailed classification on the 2-dimension space.

**Figure S8:**
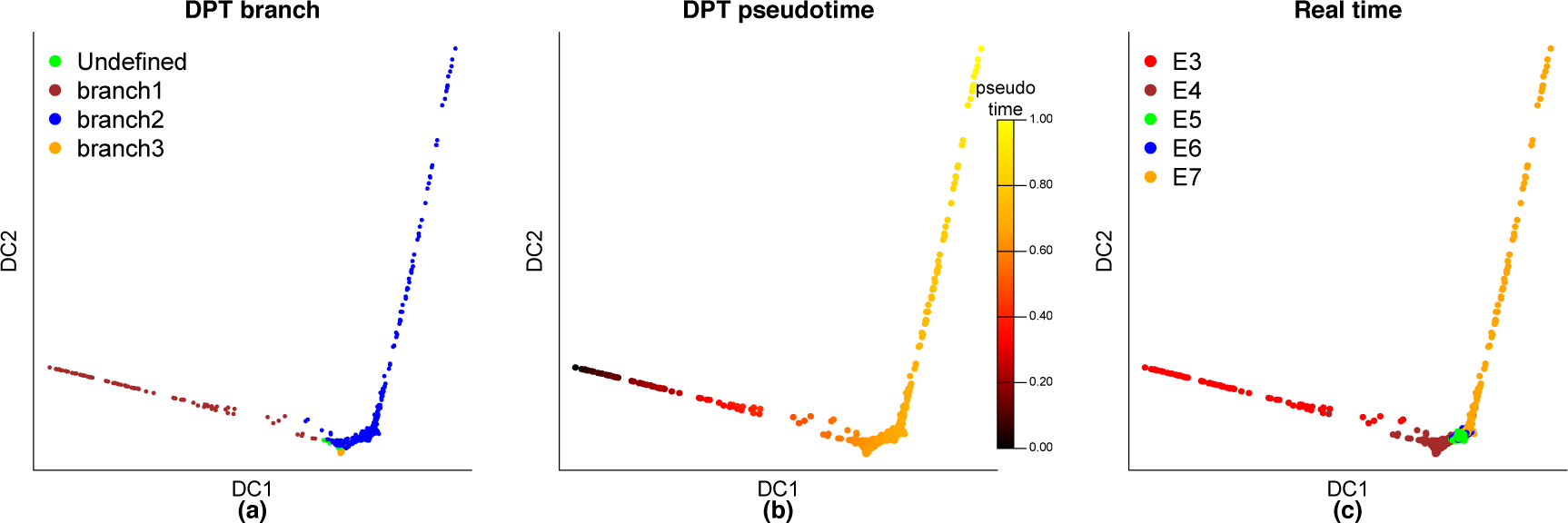
DPT analysis on HPE data. The DPT code “run a general example.m” downloaded from supplementary material of [11] was performed by setting the parameter sigma as 200. (a) DPT constructed a bifurcating trajectory. (b) DPT calculated the pseudotime of each cell. (c) The real time of each cell is displayed on the trajectory reconstructed by DPT.

**Figure S9:**
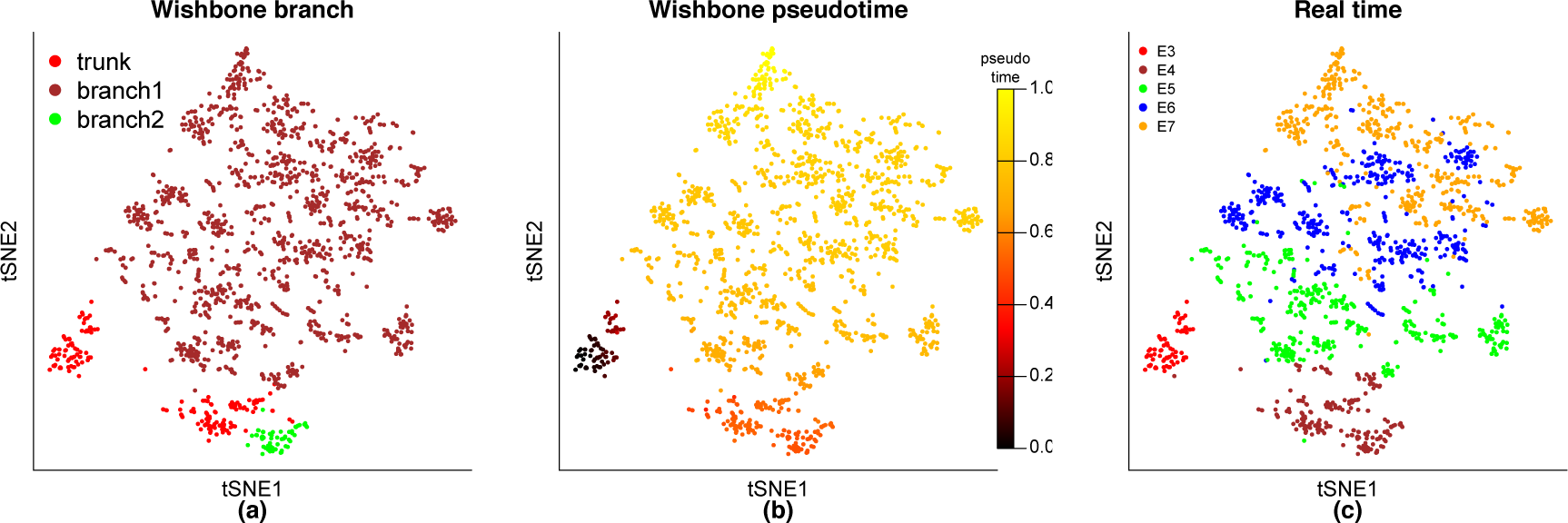
Wishbone analysis on HPE data. The Wishbone was performed by setting the parameters pca and k as 100 and 30, respectively. The visualization of results in Wishbone is demonstrated through tSNE. (a) Wishbone reconstructed a bifurcating trajectory. (b) Wishbone calculated the pseudotime of each cell. (c) The real time of each cell is displayed on the trajectory reconstructed by Wishbone.

**Figure S10:**
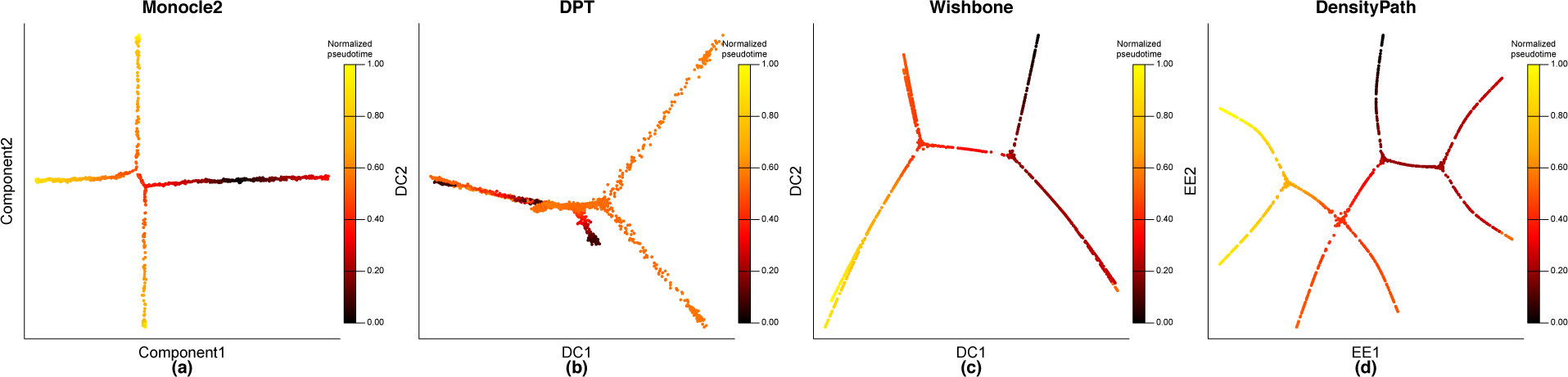
Comparison of the pseudotime reconstructed by different method on PHATE simulation data. (a) The pseudotime reconstructed by Monocle2. The Monocle2 code “analysis complex tree structure.R” was performed by setting the parameters lambda and *s* to 1000 and 0.5 respectively using DDRTree method. (b) The pseudotime reconstructed by DPT. The DPT code “run a general example.m” downloaded from supplementary material of [11] was performed by setting the parameter sigma as 150. We carried out 4 times DPT analysis: We first applied DPT to the whole cells and obtained 3 branches with the number of cells 419, 632 and 337 respectively; then we performed 3 times DPT analysis on 3 branches, separately, resulting a total of 9 branches. (c) The pseudotime reconstructed by Wishbone. The Wishbone was performed by setting the parameters pca and k as 100 and 30, respectively. (d) The pseudotime reconstructed by DensityPath.

**Figure S11:**
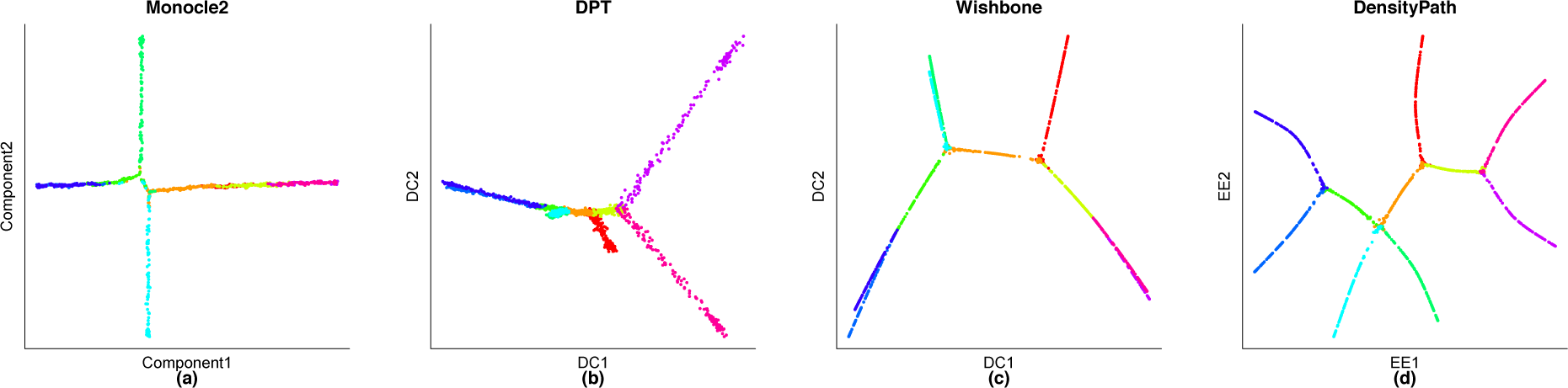
Comparison the branches assigned by different methods to the real branches assignment on PHATE simulation data. Here, the 10 colors correspond to the 10 segments of path according to the known structure. The same procedures and parameter settings for these methods were taken as Figure S10. (a) The branches assigned by Monocle2. (b) The branches assigned by DPT. (c) The branches assigned by Wishbone. (d) The branches assigned by DensityPath.

**Figure S12:**
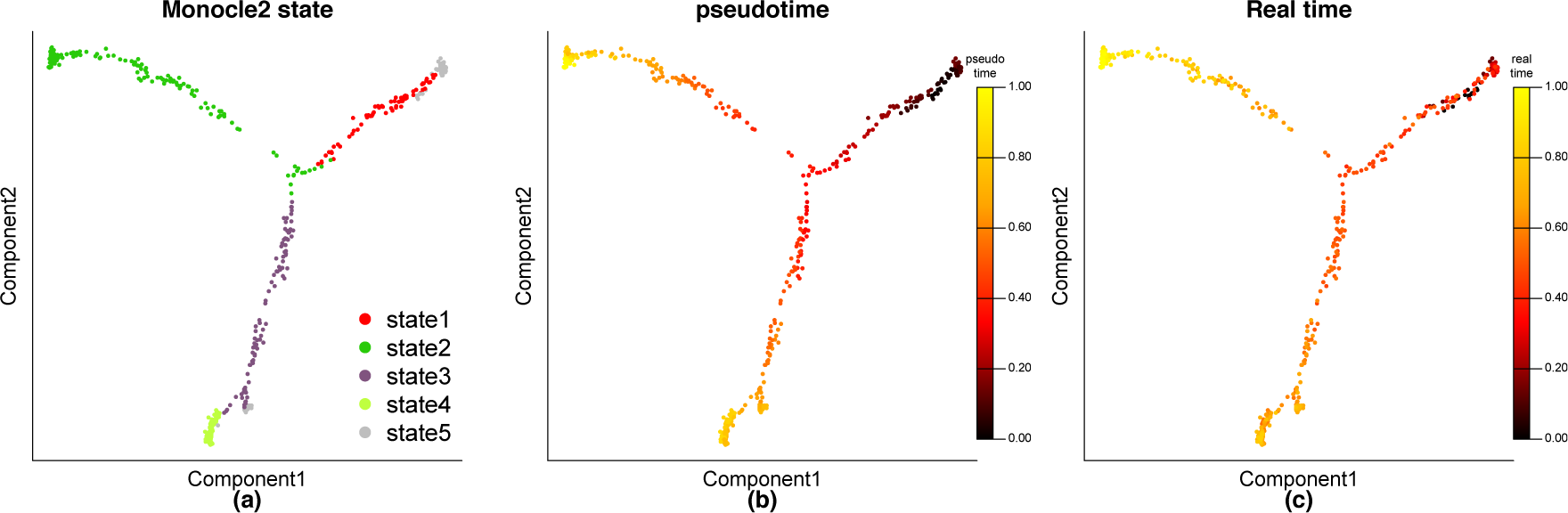
Monocle2 analysis on simulation data SLS3279. The Monocle2 code “analysis complex tree structure.R” was performed by setting the parameters lambda and *s* to 0.0464 and 0.05 respectively using DDRTree method. (a) The pseudotime reconstructed by Monocle2. (b) The pseudotime reconstructed by DPT. (c) The pseudotime reconstructed by Wishbone. (d) The pseudotime reconstructed by DensityPath.

**Figure S13:**
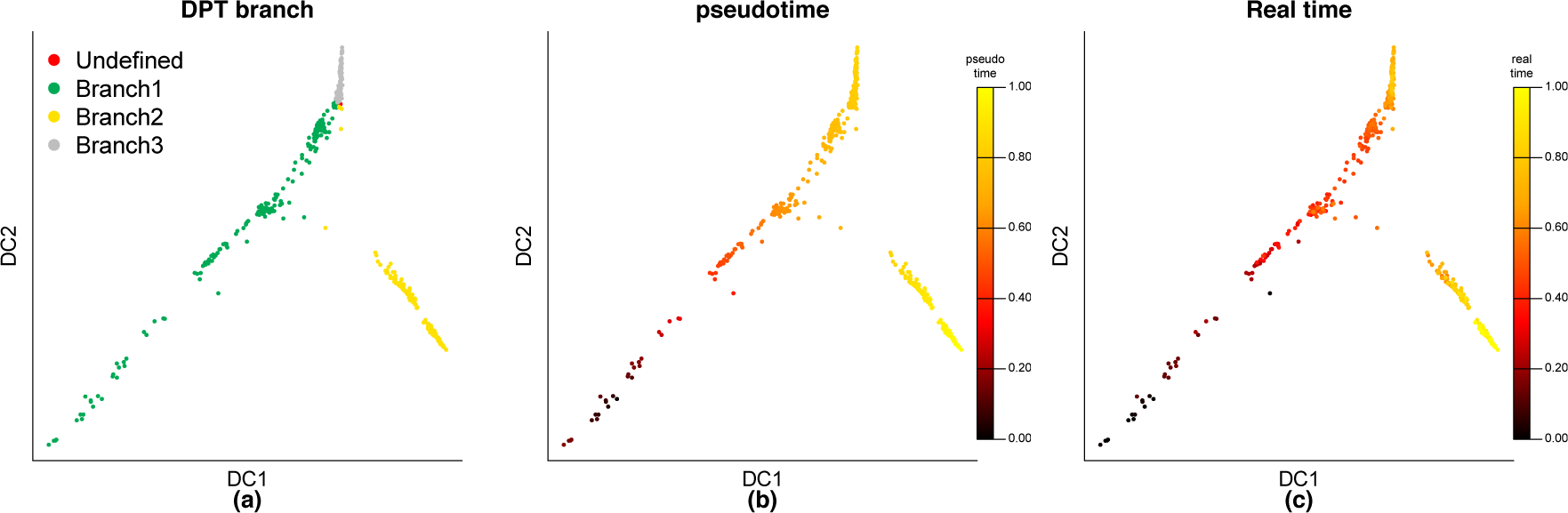
DPT analysis on simulation data SLS3279. The DPT code “run a general example.m” downloaded from supplementary material of [11] was performed by setting the parameter sigma as 2.5. (a) DPT reconstructed a bifurcating trajectory. (b) DPT calculated the pseudotime of each cell. (c) The real time of each cell is displayed on the trajectory reconstructed by DPT.

**Figure S14:**
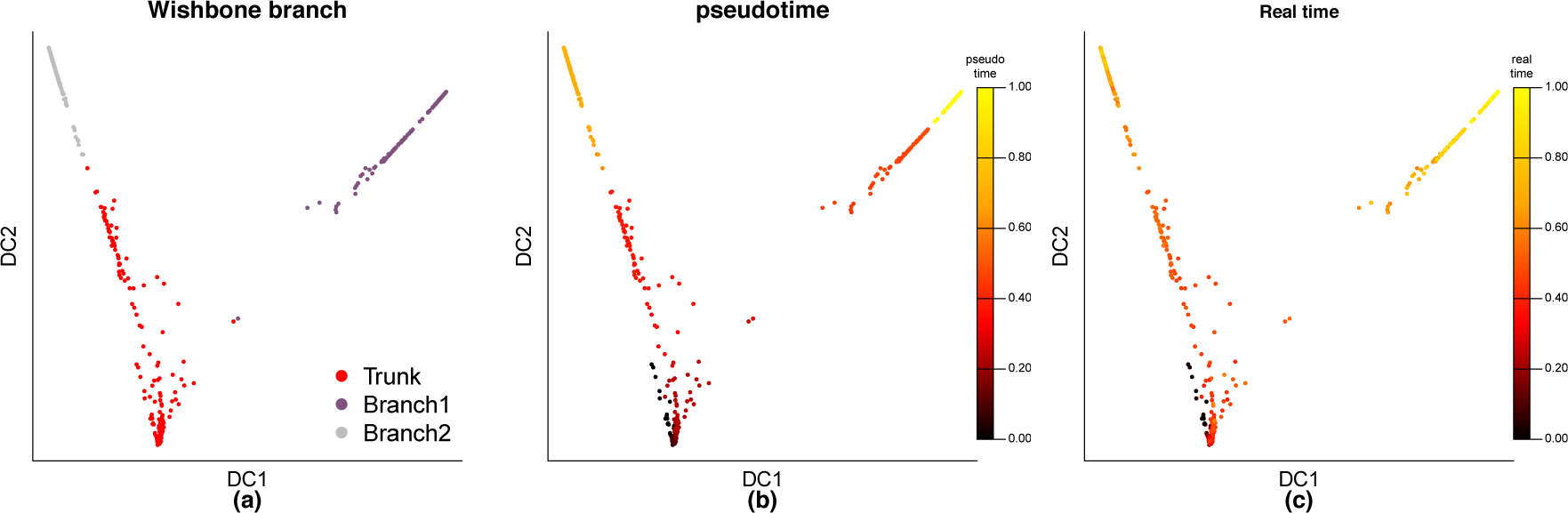
Wishbone analysis on simulation data SLS3279. The Wishbone was performed by setting the parameters pca and k as 20 and 70, respectively. (a) Wishbone reconstructed a bifurcating trajectory. (b) Wishbone calculated the pseudotime of each cell. (c) The real time of each cell is displayed on the trajectory reconstructed by Wishbone.

**Figure S15:**
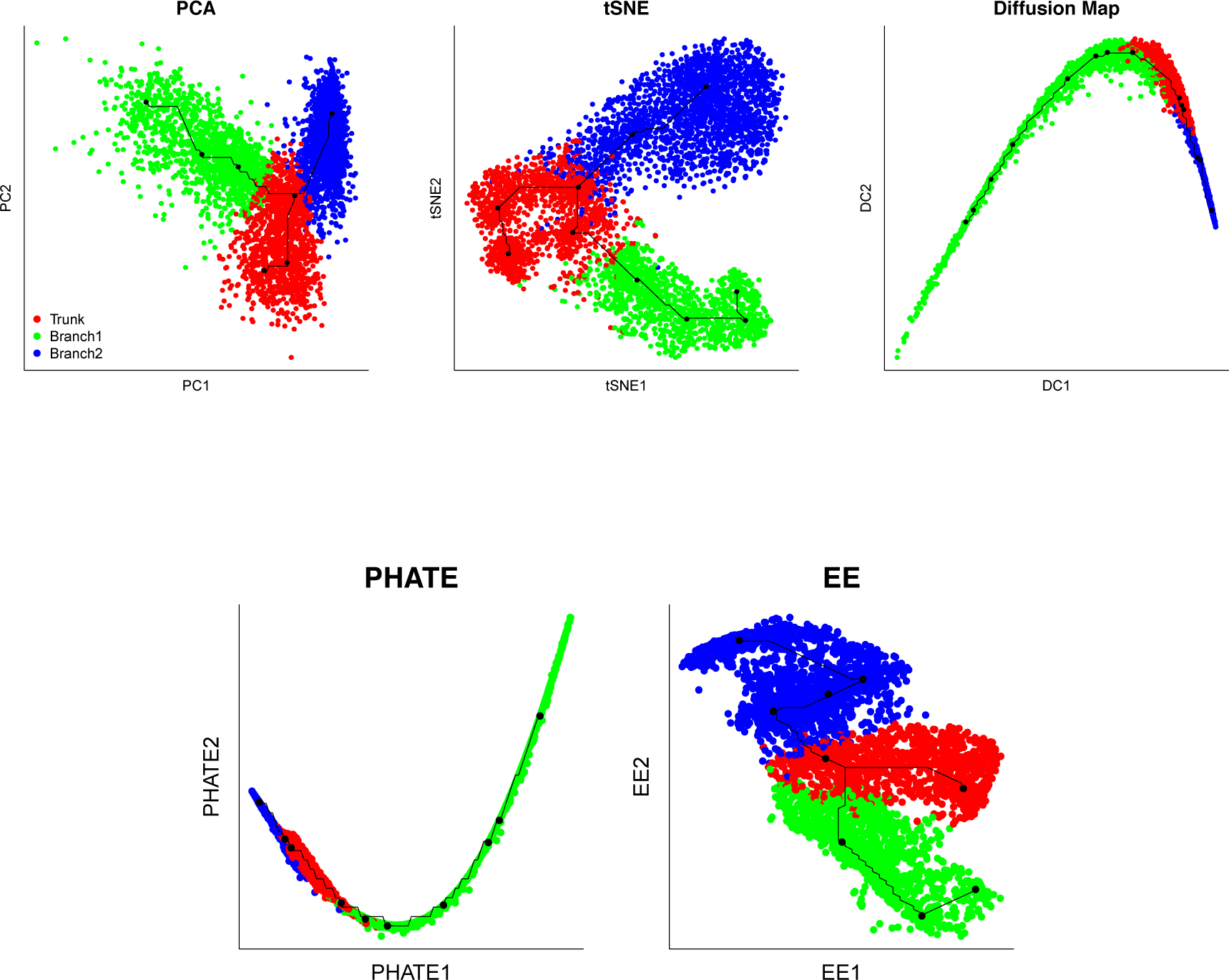
Comparison of different dimension reduction methods, PCA, tSNE, Diffusion Map, PHATE, EE on HPSCs data. Different colors reflect the assignment of cells in Wishbone while the black curves correspond to their trajectories reconstructed according to the procedures described in A2-A4 of DensityPath in Methods.

**Figure S16:**
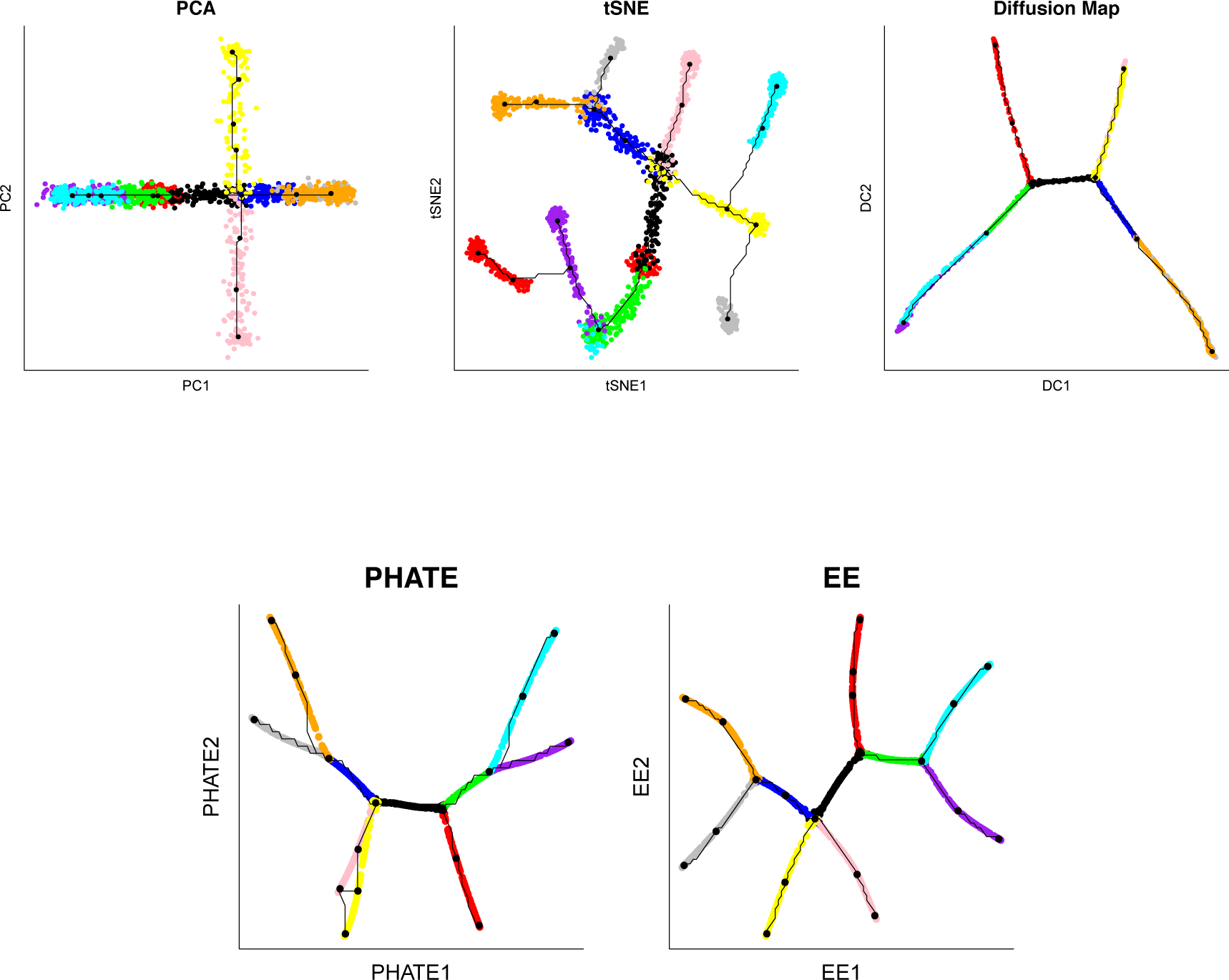
Comparison of different dimension reduction methods, PCA, tSNE, Diffusion Map, PHATE, EE on PHATE simulation data. Different colors reflect the assignment of cells in PHATE while the black curves correspond to their trajectories reconstructed according to the procedures described in A2-A4 of DensityPath in Methods.

